# GIGYF2/4EHP-Mediated Translational Attenuation Maintains Cellular Homeostasis Following Ionizing Radiation

**DOI:** 10.64898/2026.07.01.735543

**Authors:** Tom McGirr, Darrin McKenzie, Hashem Almousa, Okan Onar, Patric Harris Snell, Susanta Chatterjee, Aidan Kilmartin, Yanyu Chen, Reese Ladak, Parisa Naeli, Tamas Sessler, Sarah Maguire, Colin Adrain, Karl T. Butterworth, Alfredo Castello, Nahum Sonenberg, Robert Graham, Seyed Mehdi Jafarnejad

## Abstract

Translational regulation is a critical component of the cellular response to environmental stress. Ionizing radiation (IR) exists naturally at low doses (*e.g*., cosmic rays and radioactive materials) but is applied at much higher doses in clinical settings, where accelerated photons (X-rays) and particle beams (protons, ions etc) are utilized for the treatment of cancer. While the effects of IR on DNA damage and cell cycle are well established, its impacts on cellular RNA metabolism remains less understood. In particular, the role of the mRNA translation machinery in shaping the early cellular response to IR is largely unexplored. Here, we demonstrate that IR induces an acute and persistent translational repression. This acute repression is independent of the mTOR and Integrated Stress Response pathways, which are known regulators of mRNA translation in response to environmental cues. Instead, we discovered that the translational repression is, at least partially, mediated by the GIGYF2/4EHP translational repressor complex. We show that GIGYF2/4EHP recruitment to the mRNAs upon IR exposure is driven by rapidly enhanced interactions with RNA-binding proteins such as ZFP36 and components of the miRNA-Induced Silencing Complex (miRISC) that are poised on their target mRNAs. Importantly, the presence of the GIGYF2/4EHP complex is required for the maintenance of proteostasis and cell viability following irradiation. Together, our results establish mRNA translational control as a key determinant of cellular response to IR and identify GIGYF2/4EHP as a critical component of this adaptive mechanism.

## Introduction

In addition to its capacity for rapid regulation in response to both external and internal cues, the mRNA translation machinery is finely tuned at both global and transcript-specific levels. This enables coordinated or differential modulation of translation efficiency across large sets of mRNAs, as well as selective regulation of individual transcripts, thereby adjusting protein output in a highly dynamic manner. Such flexibility makes translational control a powerful mechanism for cellular adaptation to environmental stress (1).

The crucial role of reprogramming of translation has been recognized in response to several forms of stress such as viral infections (2), oxidative stress (3), genotoxic agents (4–7), and UV irradiation (8,9). In contrast, the role of translational regulation in the response to ionizing radiation (IR) is less well defined, as previous research primarily emphasised the transcriptional changes that underlie sustained shifts in gene expression. The studies that have characterized translational regulation mainly focused on later changes at ≥4 hours after ionizing radiation (10–12), which may confound the early impacts of IR upon translation due to indirect effects from subsequent cell cycle arrest and cell death. Ionizing radiation causes immediate damage to RNAs through direct strand breaks (13), oxidative damage (14,15), or ROS-induced protein-RNA crosslinks (16), all of which could compromise translational fidelity and proteostasis. These rapid perturbations of the RNA pool and translational machinery suggest that early translational regulation could serve as a critical acute stress-response mechanism to limit the impact of these damages.

A better understanding of how early translational regulation contributes to the cellular response to IR is important in several contexts, particularly radiotherapy-based treatments of cancers such as glioblastoma (GBM), where the efficacy of treatment remains limited due to inherent resistance (17). mRNA translation is primarily regulated at the initiation step, which in eukaryotic cells commences with the binding of eukaryotic translation initiation complex eIF4F to the m^7^G mRNA cap. eIF4F is composed of the cap binding subunit, eIF4E, the scaffolding protein eIF4G and the RNA helicase eIF4A. This complex then recruits the 43S pre-initiation complex, composed of the eIF2-GTP-Met-tRNA^iMet^ ternary complex, eIF1, eIF1A, eIF3, and the 40S ribosomal subunit, to form the 48S complex. The 48S complex scans the 5′ UTR until the start codon is recognized, prompting the joining of the 60S ribosomal subunit and formation of the translation-competent 80S ribosome. Several signalling pathways regulate the specificity and efficiency of translation at the level of initiation (18,19). Previous studies implicated two main signaling pathways in translational regulation of cellular response to IR: the integrated stress response (ISR) and mTOR. Activation of ISR kinases (GCN2, PERK, PKR, and HRI) by diverse stresses leads to phosphorylation of eIF2α, resulting in global inhibition of cap-dependent initiation, while selectively enhancing translation of stress-responsive transcription factors such as ATF4 (20,21). IR reportedly activates the ISR within 24 hours in GBM cells through PERK-dependent ER stress, highlighting its potential role in cellular response to this type of insult (22). mTOR also has been implicated in regulation of translation in irradiated HEK293T and Jurkat cells through a p53-dependent pathway downstream of double-stranded break lesions (10,11,23). mTOR primarily regulates translation through phosphorylation of 4E-BP1/2, which binds to eIF4E in its unphosphorylated state, preventing its inhibitory effects on eIF4F formation through eIF4E sequestration (21). While both mTOR and ISR have been implicated in the cellular response to radiation, the capacity of these pathways to modulate the early translational response to IR remains ill-defined.

mRNA translation could also be regulated via transcript-specific factors such as microRNAs and RNA-binding proteins (RBPs). Recent studies have demonstrated the critical role of the cap-binding eIF4E Homologue Protein (4EHP, also known as eIF4E2) in mediating the translational repression of the microRNA and RBP targeted mRNAs (24–28). 4EHP is recruited to mRNAs bound by specific RBPs or microRNA-Induced Silencing Complex (miRISC) through the scaffolding protein GRB10-interacting GYF protein 2 (GIGYF2), mostly through interactions between its GYF motif with the PPGL motif in the partner RBPs (26,29–33). 4EHP is 30% identical and 60% similar in its amino acid sequence to eIF4E. However, unlike eIF4E, 4EHP is unable to interact with eIF4G to promote translation and instead inhibit translation (34). Therefore, its binding to the cap results in repression of cap-dependent mRNA translation initiation (24). The role of 4EHP in stress-induced translational repression is being increasingly recognized, including in the microRNA directed repression of interferon response during viral infection (35,36), repression of secretory proteins during ER stress (37), and in wider translational repression following genotoxic chemotherapy treatments (7,38,39). However, its role in the response to IR is unknown.

Here, we report a rapid repression of translation following exposure of GBM cells to a clinically relevant dose of X-ray ionizing radiation. Unexpectedly, this response appears entirely independent of previously described IR-induced regulatory mTORC1 and ISR pathways and is instead mediated by a mechanism driven by 4EHP. We show that this mechanism is enabled by enhanced interactions between the GIGYF2/4EHP complex and RBPs such as ZFP36 and miRISC, which recruit it to the mRNA. Consistently, loss of this GIGYF2/4EHP-dependent repression results in increased sensitivity to ionizing radiation, implicating early translational control as a critical and previously underappreciated determinant of cellular response to ionizing radiation.

## Materials and Methods

### Cell lines and culture conditions

U251-MG, U87-MG, U343-MG, and T98G human glioblastoma cells (ATCC, Waltham, MA), and HEK293T cells (Thermo Fisher Scientific, Cat. # R70007) were grown in Dulbecco’s Modified Eagle Medium (DMEM; Gibco, Cat. # 41965039) supplemented with 10% Foetal Bovine Serum (Gibco, Cat. # 10270106), 100 U/mL penicillin and 100 µg/mL streptomycin (Gibco, Cat. # 15070063). All cell lines were tested bi-monthly for mycoplasma contamination (abm, Cat. # G238) and grown at 37°C in 5% CO_2_.

### X-ray irradiation

Irradiation exposures were performed using a 0.3 mm copper filter at a dose rate of 1.2 Gy/min using the Multirad160 (Precision X-Ray, Inc., Madison, CT, USA), calibrated according the Institute of Physics and Engineering in Medicine (IPEM) code of practice (40).

### Lentivirus shRNA packaging and generation of knockdown cell lines

HEK293T cells were transfected in 6-well plates with 1.25 µg shRNA plasmid along with equivalent amount of psPAX2 (Addgene, plasmid 12260) and pMD2.G (Addgene, plasmid 12259) in 3:1 ratio in OptiMEM (Thermo Scientific, catalog no. 51985091) using Lipofectamine 2000 (Thermo Fisher Scientific, Cat. # 11668027). 48 h and 72 h after transfection, media was collected and filtered (Filtropur S, 0.2 µm, Sarstedt, Cat. # 83.1826.001) after a brief centrifugation at 1500xg for 5 min. The collected virion-containing media was added directly to U251 cells, which were then selected in 1 µg/mL Puromycin dihydrochloride (Sigma, Cat. # P8833) for 1 week prior to validation of knockdown by western blotting. The following shRNAs were used in this study: Non-Targeting Control shRNA (shCTRL; Sigma, Cat. #SHC002), *EIF4E2* shRNA#1 (Sigma, Cat. #TRCN0000152006), *EIF4E2* shRNA#2 (Sigma, Cat. #TRCN0000280916), and *GIGYF2* shRNA (Sigma, Cat. #TRCN0000135088).

### Polysome profiling

Cells were seeded in 150 mm dishes and treated at 80% confluency, 1 h pre-treatments with DMSO (Sigma-Aldritch, Cat. # D2650) or ISRIB (Merck, Cat. # SML0843) were performed when indicated. After treatment with 100 μg/mL Cycloheximide (Sigma, Cat. # 01810) for 5 min, cells were collected by centrifugation at 4°C for 5 min and lysed in 500 μL hypotonic buffer (5 mM Tris–HCl, pH 7.5, 2.5 mM MgCl_2_ (Millipore, Cat. # M1028),1.5 mM KCl (Merck, Cat. # 529552), complete EDTA-free protease inhibitor cocktail (Roche, Cat. # 04693159001), 100 μg/mL Cycloheximide (Scientific Laboratory Supplies, Cat. # 01810), 2 mM DTT (Thermo Fischer, Cat. # R0861), 200 U/mL RiboLock RNase Inhibitor (Thermo Fisher Scientific, Cat # EO0382), 0.5% v/w Triton X-100 (Merck, Cat. # 648463), and 0.5% v/w sodium deoxycholate (Thermo Scientific Chemicals, Cat. # B20759). The lysates were cleared by centrifugation at 20,000 × g for 10 min at 4°C. 300 μg of RNA, measured by NanoDrop 2000 (Thermo Fisher Scientific), was loaded onto 10%-50% sucrose gradients, generated with sucrose solutions (Invitrogen, Cat. # 15503022) using the Gradient Master™ (BioComp, Model 108). The samples were sedimented by velocity centrifugation at 36,000 × rpm for 2 h at 4°C using SW40Ti rotor in Optima L-80XP ultracentrifuge (Beckman Coulter). Absorbance at 254 nm was measured from lower to higher sucrose gradients using an ISCO gradient fractionation system and the optical density at 254 nm was continuously recorded with a Foxy JR Fractionator (Teledyne ISCO).

### Pulse labelling with methionine analogue l-homopropargylglycine

Cells were seeded at 20,000 cells/well in a 96-well plate and treated with the indicated treatment on the following day. Click-iT™ HPG Alexa Fluor™ 488 Protein Synthesis Assay Kit (Invitrogen, Cat. # C10428) was used to measure incorporation of 50 μm l-homopropargylglycine (HPG), an alkyne-containing methionine analogue during a 30 min pulse. Cells were next fixed with 4% formaldehyde (Sigma-Aldrich, Cat # 252549) for 15 min and permeabilised using 0.5% Triton X-100 solution. Plates were incubated with Alexa Fluor 488 Azide dye in the presence of 4 mM CuSO_4_ for 30 min in the dark. Following washes, fluorescent signal was read by excitation at 488nm and absorbance at 505nm using the GloMax® Discover (Promega, Cat. # GM3000).

### PI staining and Flow Cytometry Analysis

Cells were seeded at 100,000 cells/well in 6-well plates and treated with 2 Gy the following day. After 1 h of treatment, the cells were trypsinized and pelleted by centrifugation at 2400 rpm for 5 min, resuspended in PBS +10% FBS and fixed in 4 mL of 100% ethanol overnight at 4°C. Cells were washed in PBS +10% FBS and resuspended in DNA staining solution 20 μg/ml of Propidium Iodide (Sigma, Cat. # P4864) and 200 μg/mL of RNase A (Thermo Fisher Scientific, Cat. # EN0531) in PBS for 30 min at room temperature in the dark. The stained cells were then processed on the BD Accuri™ C6 by separating the singlet population and measuring fluorescent intensity after 488 nm excitation.

### Annexin V / PI staining and Flow Cytometry Analysis

Cells were seeded at 100,000 cells/well in 6-well plates and treated with 2 Gy the following day. After 48 h of treatment, the cells were trypsinized and collected with the growth media, the adherent and detached cells were then pelleted by centrifugation at 2,400 rpm for 5 min and resuspended in PBS. The cell suspension was incubated with 5 μL of FITC-Annexin V dye (BD Pharmingen™ FITC Annexin V, Cat. #556420) for 30 min in the dark and 10 μL of Propidium Iodide stain was added immediately before analysis with the BD FACSymphony™ A1 Cell Analyzer by segregating the singlet population and measuring the fluorescent intensity after 488 nm/495 nm excitation.

### ProteoSTAT staining and Flow Cytometry Analysis

Protein aggregates were determined using the PROTEOSTAT Aggresome detection kit (Enzo Life Sciences #ENZ-51035-0025) according to the manufacturer’s instructions. Following treatment with indicated dose of radiation for 48 h, cells were collected by trypsinization, fixed in 4% formaldehyde for 30 minutes, prior to permeabilization with 0.5% Triton X-100, 3 mM EDTA (pH:8) for 30 min. Cells were then stained with a 1:7,500 dilution of ProteoSTAT aggresome detection reagent and were then processed with the BD Accuri™ C6 by segregating the singlet population and measuring the fluorescent intensity after 488 nm excitation.

### Surface sensing of translation (SUnSET) assay

Cells were seeded at 2.5 × 10^6^ cells in 60 mm well plates and pre-treated with ISRIB or DMSO for 1 h, prior to treatment with Tunicamycin (Sigma Aldritch, Cat. # T7765) for 4 h. Puromycin dihydrochloride (Sigma, Cat. # P8833) was added to the media at 2 μg/mL final concentration for 20 min prior to cell lysis with RIPA buffer (50 mM Tris–HCl pH 7.4, 150 mM NaCl (Sigma-Aldritch, Cat. # 31434), 2 mM EDTA (Merck, Cat. #324503), 1% NP-40, 0.1% SDS (Merck, Cat. #75746) supplemented with 1 mM sodium orthovanadate (WRR International, Cat. #J60191), and complete EDTA-free protease inhibitor tablet cocktail (Roche, Cat. # 11836170001)). Cells used as negative controls were pre-treated with 100 μg/mL of cycloheximide for 5 min in addition to puromycin treatment or collected without puromycin treatment. Total protein concentration was measured by Bradford protein assay (Bio-Rad, Cat. # 5000006), 10 μg of cell lysates were made up in 5x loading buffer (0.25% Bromophenol blue, 0.5 M DTT, 50% glycerol, 10% SDS, and 0.25 M Tris-Cl pH 6.8) and loaded in a 10% SDS-PAGE gel for western blotting.

### Western blotting

Cells were washed with cold PBS and scraped prior to pelleting by centrifugation at 12,000xg for 1 min. Cell pellets were lysed in RIPA buffer (as defined above). Total protein concentration was measured by Bradford protein assay and samples were made up in 5x loading buffer (as defined above). Samples were boiled for 3 min, followed by incubation on ice for 5 min. Samples were loaded onto SDS gels for electrophoresis and wet transferred onto PVDF membrane (Merck, Cat. # IPFL00010) with the BioRad Mini Trans-Blot® Cell and Criterion™ Blotter system. Membranes were blocked with 5% BSA (Thermo Fisher Scientific, Cat. # A9647) at room temperature for 1 h, incubated with the indicated primary antibodies suspended in 5% BSA overnight at 4°C and secondary antibody (1:2000) suspended in 5% BSA for 2 h at room temperature. Blots were then imaged using Syngene™ G:BOX Chemi XX9 or the Cytivia™ Amersham ImageQuant™800, and Clarity™ ECL substrate (BioRad, Cat. #1705060S). The following antibodies were used in this study: rabbit anti-phospho-RPS6 (S240/244) (Cell Signalling, Cat. # 5364), rabbit anti-RPS6 (Cell Signalling, Cat. # 2217), rabbit anti-phospho-4E-BP1 (Cell Signalling, Cat. # 2855), rabbit anti-4E-BP1 (Cell Signalling, Cat. # 9644), rabbit anti-phospho-eIF2α (Abcam, Cat. # ab32157), rabbit anti-eIF2α (Cell Signalling, Cat. # 5324), rabbit anti-PARP (Cell Signalling, Cat. # 9542), rabbit anti-Vinculin (Cell Signalling, Cat. # 13901S), mouse anti-β-actin (Sigma, Cat. # A5441), mouse anti-puromycin (Merck, Cat. # MABE343), rabbit anti-eIF4E2 (GenTex, Cat. # GTX825224), rabbit anti-TNRC6C (Bethyl, Cat #A303-969A), rabbit anti-GIGYF2 (Proteintech, Cat. # 24790-1-AP), rabbit anti-ZFP36 (Proteintech, Cat. # 12737-1-AP), rabbit anti-ATF-4(Cell Signalling, Cat. #D4D8, rabbit anti-CHOP (Cell Signalling, Cat. #D46F1), anti-rabbit IgG HRP-linked (Thermo Fisher Scientific, Cat. # 7074S), anti-mouse IgG HRP-linked (Thermo Fisher Scientific, Cat. # 7076S). The uncropped images of all blots are provided in Supp. Fig. 6-12.

### Colony formation assay

U251 cells were seeded into 6-well plates at different densities depending on the dose of radiation delivered the following day (0 and 2 Gy: 2000 cells, 4 Gy: 4000 cells, 4 Gy: 6000 cells and 8 Gy: 8000 cells) and 1 h pre-treatments with DMSO or ISRIB were performed where indicated. After 10 days, cells were rinsed with PBS before staining with colony fixation-staining solution (Crystal Violet (Sigma-Aldrich, Cat. # C0775), w/v in 95% Methanol) for 15 min, followed by two brief washes in water and drying overnight. Afterwards, the number of colonies (diameters > 50 μm) were counted with the Oxford Optronix GelCount™ system (v1.1.2.0; Oxford Optronix). Plating efficiencies were calculated from the raw counts, and the surviving fraction was calculated as the proportion of colonies compared to control cells.

### Incucyte proliferation assay

Cells were plated at 2,000 cells/well in 96 well plates prior to treatment on the following day. Cells were placed in the Sartorius Live-Cell Imaging and Analysis Instrument: Incucyte® SX5, imaged every 6 h for a week and images were analysed to calculate the percentage confluency normalized to the starting percentage confluency.

### RNA-Interactome Capture (RIC) assay

150 mm plates were seeded with 15 × 10^6^ for 1 h treatment or 10 × 10^6^ for 24 h treatment prior to treatment the following day. After the desired time, cells were washed in PBS and dry plates were either mock UV irradiated or crosslinked with 150 mJ/cm^2^ at 254 nm using the Analytik Jena CL-3000 on ice. Cells were immediately lysed in lysis buffer (20 mM Tris-HCl pH 7.5, 500 nM LiCl (Merck, Cat. #62476), 0.5% LiDS (Carl Roth, Cat. # CN25.3), 1 mM DTT (Thermo Fischer, Cat. # R0861), 1 mM EDTA (Merck, Cat. #324503)) and cleared by sequential passages through 27G and 23G needles. 300 μL/sample of Dynabeads™ Oligo(dT)_25_ beads (Invitrogen™, Cat. #61002) were pre-washed with lysis buffer and added to cleared lysates for rotation at 4°C for 1 h. Beads were washed with 5 min incubation steps at 4°C once in lysis buffer, twice in wash buffer 1 (20 mM Tris-HCl pH 7.5, 500 nM LiCl, 0.1% LiDS, 5 mM DTT, 1 mM EDTA), twice in wash buffer 2 (20 mM Tris-HCl pH 7.5, 500 nM LiCl, 5 mM DTT, 1 mM EDTA, 0.1% DDM; Thermo Fisher, Cat. #D4641). Finally, beads were washed once with wash buffer 3 (20 mM Tris-HCl pH 7.5, 200 nM LiCl, 5 mM DTT, 1 mM EDTA, 0.1% DDM), prior to elution in 220 μL elution buffer (20 mM Tris-HCl pH 7.5, 1 mM EDTA) at 55°C. RNA concentration was measured using Nanodrop™ ND 1000 Spectrophotometer and samples were only used if 260/280 nm absorbance ratio was below 1.9 and 260/230 absorbance ratio was below 2. Eluates were then treated with 100 U of RNase A (Thermo Fisher, Cat. #R1253) and 100-400 U of RNase T1 (Sigma-Aldrich, Cat. #R1003) for 1 h at 37°C and 15 min at 50°C before storage at -80°C. 20 μL of eluate was taken to perform silver staining with SilverQuest™ Silver Staining Kit (Thermo Fisher Cat. #LC6070) per manufacturers’ protocol. For proteomic analysis, samples were processed using the protein aggregation capture (PAC) method (41). Proteins were reduced with 10 mM DTT for 30 min at room temperature, followed by alkylation with 50 mM iodoacetamide (IAA) for 30 min at room temperature in the dark. Samples were then split into aliquots of 300 μL, and 5 μL of MagResyn Hydroxyl beads (Cat #MR-HYX002) were added to each tube. Acetonitrile (ACN) was added to a final concentration of 80% (v/v), and samples were incubated for 30 min at room temperature. Beads were washed twice with 100% ACN and three times with 70% ethanol to remove residual detergent and non-specific binders. Proteins were digested overnight at 37°C with 1 μg trypsin (Promega, Cat. #V5280). The resulting peptides were dried down in a Speedvac concentrator (Eppendorf, U.K.) prior to desalting.

### Co-immunoprecipitation

Cells were washed with cold PBS and collected by scraping in lysis buffer (40 mM HEPES pH 7.5 (Sigma-Aldrich, Cat #H0887), 120 mM NaCl, 1 mM EDTA, 0.3% CHAPS (Sigma-Aldrich, Cat #331717), supplemented with complete EDTA-free protease inhibitor cocktail). Dynabeads™ Protein G (Invitrogen, Cat #10003D) were blocked with 2% BSA in PBST for 1 h and washed 3x for 2 min each in TBST. 6mg of the pre-cleared lysates were incubated with 3 µg anti-GIGYF2 antibody (Proteintech, Cat. # 24790-1-AP), 6 µg of eIF4E2 antibody (GenTex, Cat. #GTX825224), or matched amounts of IgG isotype control (Proteintech, Cat #ab172730) for 30 min in presence of RNase A (Thermo Fisher Scientific, Cat. # EN0531). 60 µL of blocked and prewashed Dynabeads™ Protein G was then added prior to incubation at 4°C overnight. Beads were washed thrice for 10 min each with wash buffer (50 mM HEPES pH 7.5, 150 mM NaCl, 1 mM EDTA, 0.3% CHAPS, supplemented with complete EDTA-free protease inhibitor cocktail). 1/6^th^ of washed beads (equivalent to 1 mg of input) was eluted in lysis buffer in 5x loading dye (as defined above) by heating at 95°C for 10 min, while 5/6^th^ (equivalent to 5 mg of input) was flash frozen in wash buffer for mass spectrometry. For mass spectrometry, beads were resuspended in 20 mM of triethylammonium bicarbonate (TEAB) buffer and proteins were reduced with 10 mM dithiothreitol (DTT) for 30 min at room temperature, followed by alkylation with 50 mM iodoacetamide (IAA) for 30 min at room temperature in the dark. Proteins were digested off the beads with 200 ng Trypsin (Promega, Cat. #V5280) overnight at 37°C. The resulting peptides were dried down in a Speedvac concentrator (Eppendorf, U.K.) prior to desalting.

### LC-MS/MS data acquisition

Peptides were desalted using in-house made STAGE (STop And Go Extraction) tips following established protocols (42). Precipitated peptides were resuspended in 20 μL of 0.1% formic acid and loaded onto C18 columns that had been pre-equilibrated with 0.1% formic acid following sequential washes with 80% methanol and 80% ACN. The columns were subsequently washed twice with 0.1% formic acid, and bound peptides were eluted using a solution of 80% ACN containing 0.1% formic acid. Eluates were dried down in a Speedvac concentrator (Eppendorf, U.K.) and resuspended in 10 µl 0.1% formic acid on the day of mass spectrometry analysis.

Liquid chromatography tandem mass spectrometry (LC-MS) analysis was performed using a Bruker TimsTOF HT mass spectrometer coupled to a nanoElute 2 nanoLC system via a CaptiveSpray ionisation source. Samples were injected directly onto a self-packed reverse phase C18 column (75 µm inner diameter, 1.5 µm particle size, Dr Maisch beads; MSWIL, Cat. # ra115.9e.0001). Buffer A comprised of 0.1% formic acid in water, while buffer B consisted of 0.1% formic acid in ACN. Peptide separation was carried out using a 71 min linear gradient at a flow rate of 250 nL/min and a column temperature of 50°C, progressing from 3% to 15% buffer A over 40 min, 15% to 25% over 65 min and 25% to 33% over 71 min. Column washing performed with 95% buffer B for 4 minutes. Data was acquired using dia-PASEF (data-independent acquisition combined with parallel accumulation-serial fragmentation) as previously described (43). Isolation window placement was optimised using py-diAID tool, resulting in a method comprising 32 MS/MS windows spanning m/z 261.1-1399.7 with a total cycle time of 1.8 seconds (44). The TimsTOF ion mobility range was set from 1/k0 = 0.57 to 1.47 Vs cm^2^, with accumulation and ramp times both set to 100 m/s.

### Data analysis

Raw data files from RIC and Co-IP experiments were analysed using Spectronaut® v19 (https://biognosys.com/software/spectronaut/) via DirectDIA. Parameters included Trypsin as the enzyme, allowing a maximum of two missed cleavages Carbamidomethylation of cysteine was defined as a fixed modification, whereas methionine oxidation and N-terminal acetylation were included as variable modifications. Data were searched against the *Homo sapiens* reference proteome (UniProt ID: UP000005640) with a peptide-level False Discovery Rate (FDR)<1%. Data analysis of the Protein Quantity file from Spectronaut was performed in R. Rows were filtered to remove any with >5 NA values in each condition for RNA-interactome capture, and >2 N/A values for AP-MS. Data were median normalized and Perseus-style minimum value imputation was performed by drawing random numbers from a normal distribution with a downshift of 1.8 standard deviations and a width of 0.3, applied independently to each sample. Principal component analyses were performed using the base R. package on log2-transformed values. For RNA-interactome capture, batch correction was performed using the ComBat R package (45). For 4EHP AP-MS, variation between matched independent replicates was controlled by incorporating replicate identity as a blocking factor in a paired limma analysis. Fold-changes and adjusted-*p*-values were calculated using the limma R. package and Benjamini-Hochberg multiple-testing correction (46). Gene ontology (GO) analysis was performed using the org.Hs.eg.db, enrichplot and clusterprofiler R. packages (47–49).

### Statistical analyses

Statistical analyses were performed using Prism 6 (GraphPad). Error bars represent standard deviation (SD) from the mean of at least three independent replicates collected on separate days and individual data points are depicted in bar graphs. Statistical significance was set a priori at 0.05. Only one observation per sample was collected.

## Results

### Ionizing radiation induces rapid, persistent translational repression in GBM cells

We first assessed the translational response of GBM cells following exposure to 2 Gy of X-ray photons; the common modality and fractionated dose of IR used in radiotherapy for glioblastoma (GBM) (50). This dose was also chosen as it doesn’t trigger cell death, as indicated by the absence of PARP cleavage or significant cell growth arrest (**Supp. Fig. 1A & B**) within the first 24 hours post-treatment, thus avoiding the confounding effects of reduced cell viability on mRNA translation (11). To assess the effect of IR on general mRNA translation, we used polysome profiling, which measures RNA absorbance across a sucrose gradient where mRNAs are sedimented based on number of associated ribosomes (51). Efficiently translated mRNAs, which are loaded with a higher number of ribosomes (polysomes) sediment in the heavier sucrose fractions, while poorly translated mRNAs sediment in the lighter fractions (sub-polysome). We observed that irradiation of U251 GBM cells resulted in a significant shift (20±1%, p<0.001) from the efficiently translated polysome to poorly translated sub-polysome fractions within 1 h post-treatment (**Fig. 1A & B**). Notably, this reduction persisted up to 48 hours post-treatment (19±1%, p<0.001; **Fig. 1B & Supp. Fig. 1C-F**). A similar effect was also observed in U87 GBM cells (**Fig. 1C & D, Supp. Fig. 1G & H**). These data suggest a rapid and sustained repression of mRNA translation in these GBM cells following treatment with a well-tolerated dose of IR.

**Figure 1:**
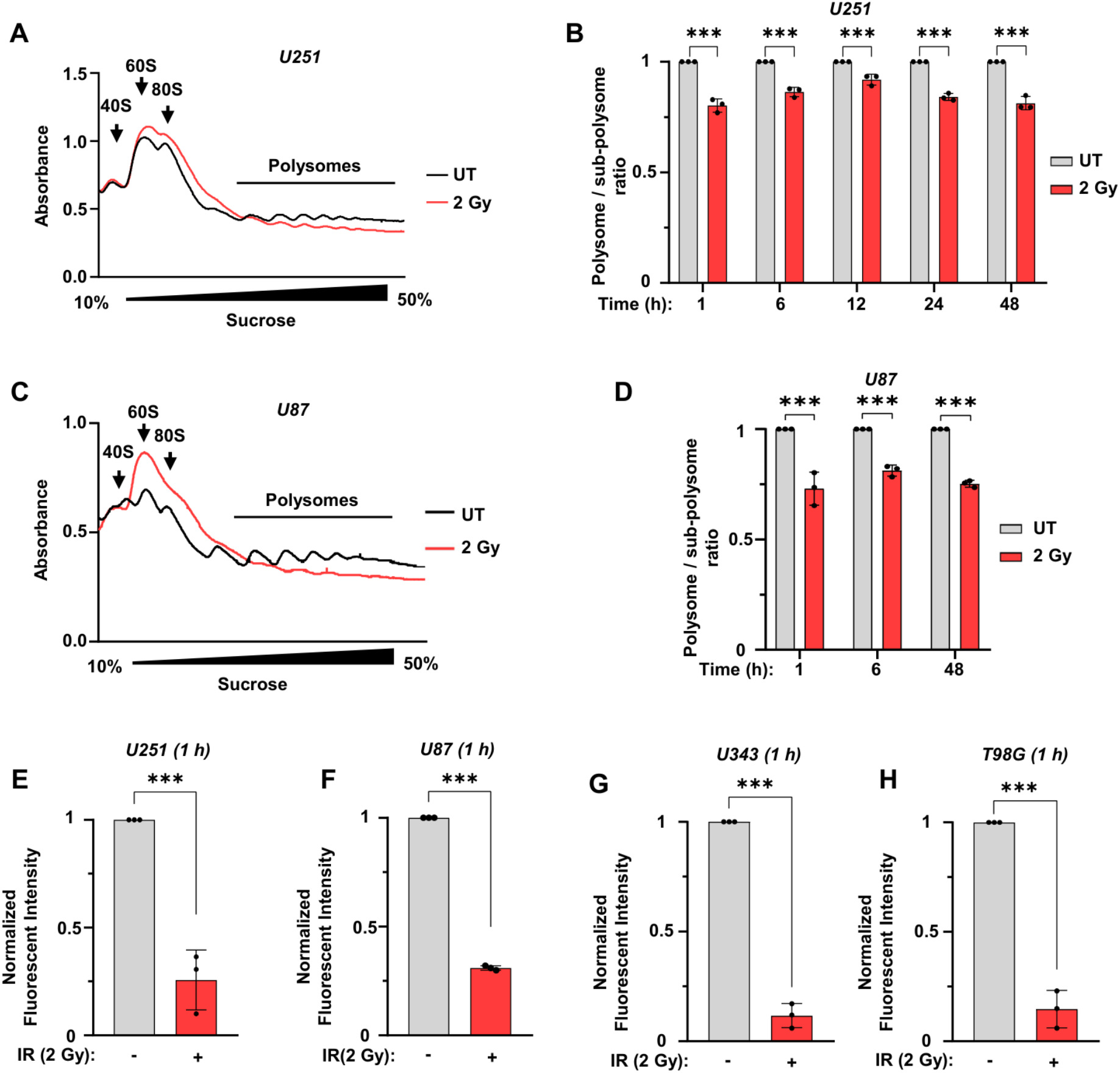
Ionizing radiation triggers rapid and sustained translational repression in GBM cells. (**A**) Representative polysome profiling traces from U251 cells 1 h following 2 Gy irradiation. (**B**) Quantification of the polysome to sub-polysome ratio derived from polysome profiling experiments in U251 cells at 1 h, 6 h, 12 h, 24 h, and 48 h following 2 Gy irradiation, normalized to time-matched untreated controls. (**C**) Representative polysome profiling traces from U87 cells 1 h following 2 Gy irradiation. (**D**) Quantification of the polysome to sub-polysome ratio in U87 cells at 1 h, 6 h, and 48 h following 2 Gy irradiation, normalized to time-matched untreated controls. (**E-H**) Quantification of nascent protein synthesis measured by metabolic pulse labelling with the methionine analogue L-homopropargylglycine (HPG) 1 h after 2 Gy irradiation. Normalized fluorescence intensity was calculated by deduction of the unstained fluorescence followed by division by untreated fluorescence in U251 (E), U87 (F), U343 (G), and T98G cells (H). Data are presented as mean ± SD; n=3, one-way Anova; ***p<0.001.

To capture the acute responses to radiation, we investigated the 1-hour post-treatment window where no coordinated cell cycle arrest was observed (**Supp. Fig. 1I & J**). We corroborated the acute reduction in the polysome/sub-polysome ratio at 1 hour post-irradiation using a pulse-labelling assay with the azide-containing methionine analogue L-homopropargylglycine for analysis of nascent protein synthesis detected following click reaction with alkyne-fluorophore (52). The treatment caused a significant reduction in fluorescent signal, denoting reduced *de novo* protein synthesis, by 1 hour after irradiation in four different GBM cell lines; U251 (74±8% reduction, p<0.001; **Fig. 1E**), U87 (69±1% reduction, p<0.001; **Fig. 1F**), U343 (88±3% reduction, p<0.001; **Fig. 1G**) and T98G (85±5% reduction, p<0.001; **Fig. 1H**). Notably, the reduction in the *de novo* protein synthesis in U251 cells strongly correlated with the dose of IR exposure, as evident by the lack of change in the fluorescent signal in cells treated with 0.1 Gy (**Supp. Fig. 1I**) and the progressively increased reduction in cells treated with 1 Gy (47±9% reduction, p<0.05; **Supp. Fig. 1J**), 2 Gy (74±8% reduction, p<0.001; **Fig. 1E**), and 6 Gy (87±3% reduction, p<0.001; **Supp. Fig. 1K**). Collectively, these data reveal a rapid repression of translation following irradiation. Critically, the onset of repression occurs before detectable cell cycle arrest or apoptosis (**Supp. Fig. 1L & M**), consistent with a mechanism that acts directly at the level of translation.

### Acute IR-induced translational repression is independent of the mTORC1 and ISR signalling pathways

To explore the underlying mechanisms of the early translational repression upon IR exposure, we examined the potential role of two key signalling pathways, namely mTORC1 and the Integrated Stress Response (ISR) pathways (53), which were previously implicated in later phases (>24 h) of IR-induced translational regulation (10–12,22). Notably, we did not observe a significant reduction in 4E-BP1 phosphorylation at Thr37/46, a marker of mTORC1 activity and an effector of mTOR-mediated translational regulation, within the first 12 hours post-IR treatment in either U251 or U87 cells (p>0.05; **Fig. 2A & B**) (54). The lack of reduced mTORC1 activity after IR was corroborated by assessing another marker of mTORC1 activity, phosphorylation of the ribosomal protein S6 at S240/S244 (p>0.05; **Supp. Fig. 2A-B**) (55). The reduced mRNA translation, despite the lack of change (in U251 cells) or increased mTORC1 activity (in U87 cells), indicate that the rapid reduction in the mRNA translation upon IR treatment cannot be explained by reduction in mTORC1 activity.

**Figure 2:**
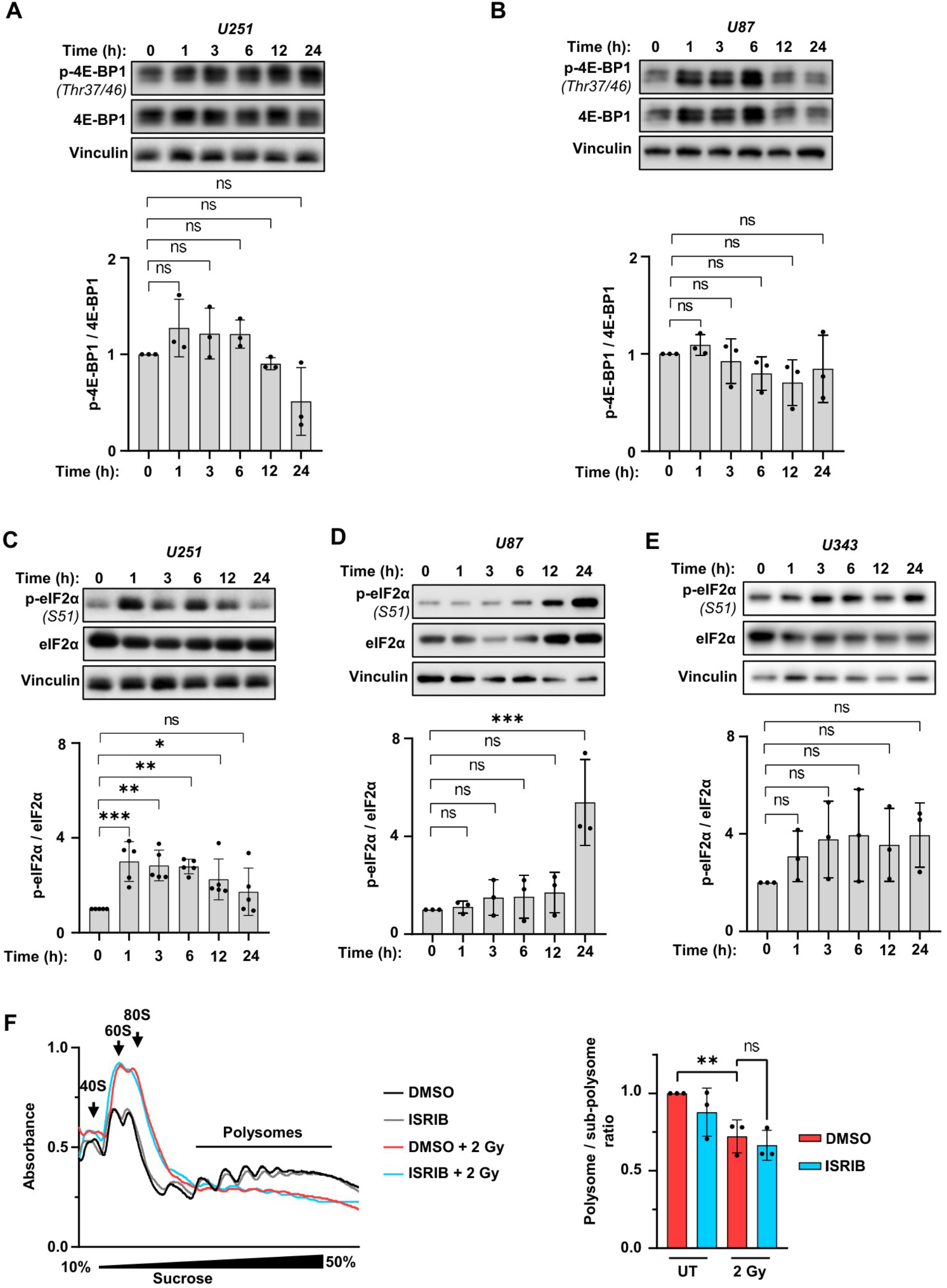
Early radiation-induced translational repression is independent of mTOR and ISR signaling. (**A & B**) *Top:* Western blot analysis of phosphorylation of 4E-BP1 (Threonine 37/46), a marker of mTORC1 activity in U251 (A) and U87 (B) cells following 2 Gy irradiation. Vinculin was used as a loading control. *Bottom:* Densitometry quantifications of p-4E-BP1/4E-BP1 in the respective cells. (**C-E**) *Top:* Western blot analysis of phosphorylation of eIF2α (Serine 51), a marker of ISR activity in U251 (C), U87 (D), and U343 (E) cells following 2 Gy irradiation. *Bottom:* Densitometry quantifications of p-eIF2α/eIF2α in the respective cells. (**F**) *Left:* Polysome profiling traces from U251 cells pre-treated for 1 h with ISRIB (100 nM) or DMSO prior to 2 Gy irradiation for 1 h. *Right:* Quantification of the polysome to sub-polysome ratio in (F). Data are presented as mean ± SD; n=3 (n=5 in C), one-way Anova; ns=non-significant, *p<0.05, **p<0.01, ***p<0.001.

We next investigated the activity of the ISR pathway after irradiation by measuring phosphorylation of eIF2α at Ser51. IR induced a transient increase in eIF2α Ser51 phosphorylation in U251 cells, peaking at 1 h post-treatment (3±0.4 fold increase, p<0.001; **Fig. 2C**) and declining by 12 hours (2.3±0.4 fold increase, p<0.05; **Fig. 2C**). In contrast, U87 cells exhibited delayed ISR activation, with a significant increase in eIF2α Ser51 phosphorylation being detected only at 24 h post-treatment (5.4±0.8 fold increase, p<0.001; **Fig. 2D**). Notably, we detected no change in eIF2α phosphorylation following irradiation of U343 cells (p>0.05; **Fig. 2E**). Despite comparable translational repression 1 hour after irradiation in these three cell lines (**Fig. 1**), eIF2α phosphorylation was not uniformly increased at 1 hour time-point across different cell lines, indicating that ISR induction may contribute to but cannot be the driver of this early radiation-induced translational repression across the GBM cells tested in this study. However, considering the significant early upregulation of ISR in the U251 cells (**Fig. 2C**), we sought to further investigate the contribution of the ISR to early translational repression in this cell line. We pharmacologically inhibited ISR signaling using ISRIB, which restores translation downstream of eIF2α phosphorylation by stabilizing eIF2B into an active conformation (56,57). Dose-response analysis identified ≥100 nM ISRIB as sufficient to rescue protein synthesis from tunicamycin-induced ISR activation in U251 cells, as assessed by SUnSET assay (**Supp. Fig. 2C**) (58). Therefore, we employed this dose in combination with IR to assess whether ISR inhibition could reverse early radiation-induced translational repression. However, pre-treatment with ISRIB had no effect on the acute IR-induced reduction of mRNA translation in U251 cells, as evidenced by lack of significant change in the polysome/sub-polysome ratio between ISRIB and vehicle-treated cells (p>0.05; **Fig. 2F**). Consistently, pre-treatment with different doses of ISRIB (10nM, 50nM, and 100 nM) had no significant effect on the fractional survival of U251 cells after irradiation, as measured by clonogenicity (p>0.05; **Supp Fig. 2D**). The lack of ISRIB-induced reversal of the translational repression in U251 cells, along with the lack of correlation between ISR activation and early repression in U87 and U343 cells suggest that ISR is not a critical driver of the IR-induced translational downregulation across GBM cells.

Together, these data suggest the presence of alternative mechanism(s) of translational regulation, besides mTORC1 and ISR, that may contribute to early translational repression in GBM cells.

### GIGYF2/4EHP mediates the rapid translational repression induced by IR

Recent studies documented a key role for the cap-binding and translational repressor protein 4EHP and its interacting partner GIGYF2 (**Fig. 3A**), in cellular response to environmental stress that affect mRNA translation such as viral infections, ribosome collisions, and ER stress (7,30,35,37,59–61). Therefore, we next investigated the roles of 4EHP and GIGYF2 in the rapid downregulation of mRNA translation in response to IR. We generated U251 cells that stably expressed two independent shRNAs against 4EHP (sh4EHP-1 and sh4EHP-2; **Supp. Fig. 3A**) or against GIGYF2 (shGIGYF2; **Supp. Fig. 3B**), and U87 cells that stably expressed sh4EHP-1 (**Supp. Fig. 3C**). Polysome profiling revealed a ∼50% reversal (p<0.001; **Fig. 3B**) of the IR-induced reduction in the polysome/sub-polysome ratio in the 4EHP knockdown cells (sh4EHP-1; 86±6) compared with the control cells (shCTRL; 71±6%). Pulse labelling assay with the methionine analogue L-homopropargylglycine further validated this observation. Accordingly, while IR treatment dramatically reduced the rate of new protein synthesis in the shCTRL cells 1 h post-treatment, 4EHP knockdown significantly reversed this reduction (25±6% reversal for sh4EHP-1, p<0.01, and 41±4% reversal for sh4EHP-2, p<0.001; **Fig. 3C**). Similarly, 4EHP knockdown in U87 cells resulted in 25±6% reversal (p<0.05; **Supp. Fig. 3D**) of translational repression 1 h post IR exposure. Furthermore, GIGYF2 knockdown resulted in a reversal (34±12%, p<0.05; **Fig. 3D**) of the IR-induced repression of new protein synthesis in U251 cells.

**Figure 3.**
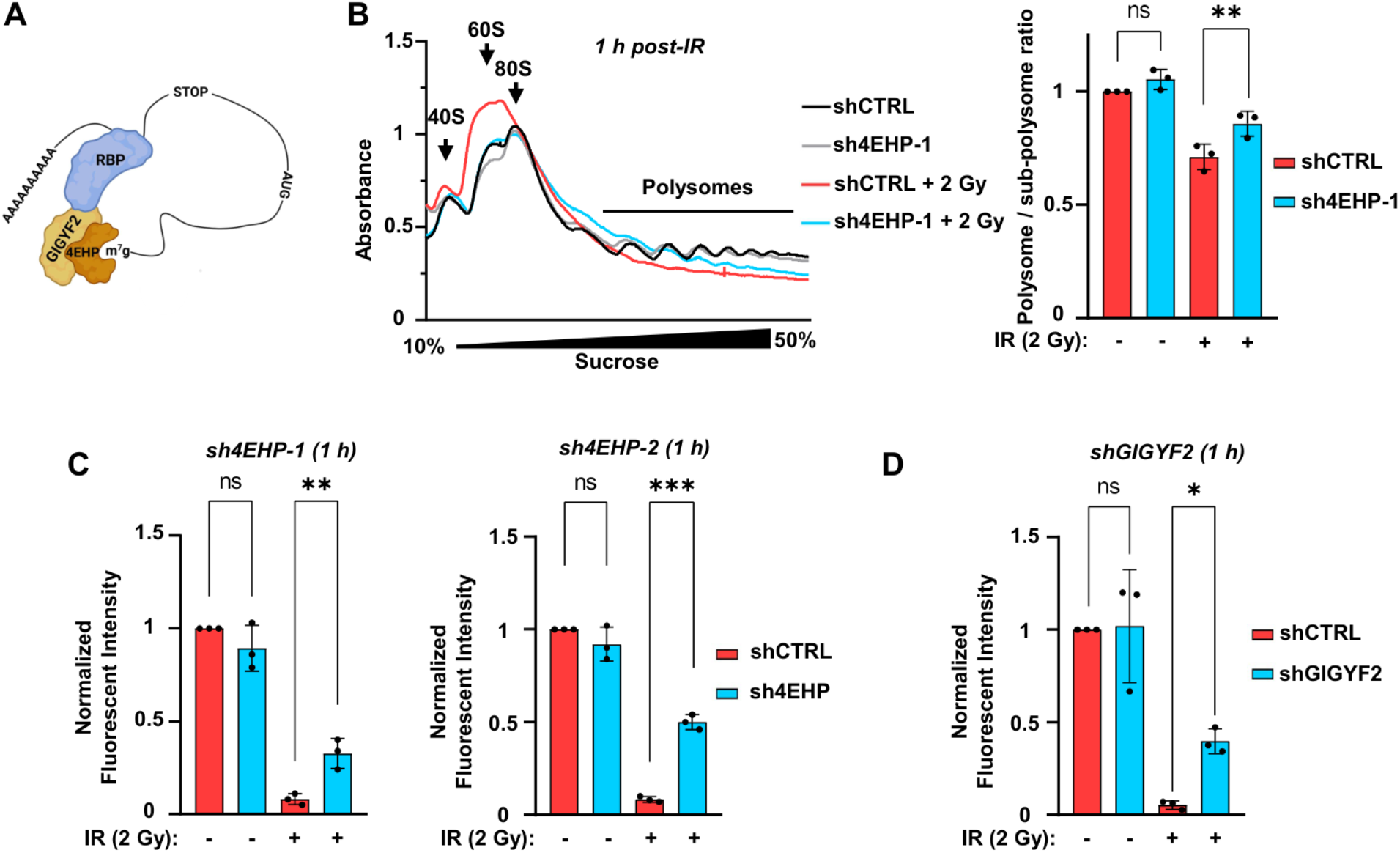
Depletion of 4EHP and GIGYF2 reverses the translational repression upon IR exposure. (**A**) Schematic model depicting the transcript-specific repression of mRNA translation by the RBP/GIGYF2/4EHP complex. Several RBPs including ZFP36, ZNF598, and the microRNA-Induced Silencing Complex (miRISC) are known to participate in this complex to repress their target mRNAs. (**B**) *Left*: Representative polysome profiling traces from the control (shCTRL) and 4EHP-knockdown (sh4EHP) U251 cells 1 h post-treatment with 2 Gy radiation. *Right*: Quantification of the polysome to sub-polysome ratios. (**C & D**) Quantification of nascent protein synthesis in shCTRL, sh4EHP-1, and sh4EHP-2 (C) and GIGYF2-knockdown (D) U251 cells measured by metabolic pulse labelling with the methionine analogue L-homopropargylglycine (HPG), 1 h post-treatment with 2 Gy radiation. Signals were normalized against the unstained samples for each group. Data are presented as mean ± SD; n=3, one-way Anova; ns=non-significant, *p<0.05, **p<0.01, ***p<0.001.

These data suggest that 4EHP and GIGYF2 are involved in the rapid repression of mRNA translation in response to IR treatment.

### GIGYF2/4EHP depletion impairs proteostasis and cellular viability upon IR exposure

We next assessed whether abrogating the GIGYF2/4EHP-mediated translational repression could negatively affect cellular homeostasis. We reasoned that enhanced mRNA translation in the absence of GIGYF2/4EHP might result in elevated levels of aberrant translation, leading to the accumulation of misfolded or otherwise abnormal translation products and subsequent activation of proteotoxic stress responses. Consistent with this hypothesis, we observed that 4EHP knockdown markedly increased the expression of ATF4 and CHOP, two well-established markers of the integrated stress response and proteotoxic ER stress 48 h upon IR exposure (2.2±0.7 and 3.3±0.3 increase for ATF4 and CHOP, respectively, p<0.05, p<0.001; **Fig. 4A**) with increased CHOP expression persisting to up to 72 h post-exposure (2.6±0.3 increase, p<0.001; **Fig. 4A**) (20). We utilized the ProteoSTAT assay, which quantifies protein aggregates through a fluorophore-conjugated antibody. The assay’s sensitivity to aggregated proteins was validated by treatment with the proteasome inhibitor MG132, which induced robust aggregate accumulation, where we noted a 2-fold increase in red fluorescent intensity **(Supp. Fig. 4A)**. ProteoSTAT revealed a 1.6-fold increased aggregation in 4EHP knockdown compared with the control cells after 2 Gy radiation (p<0.01; **Fig. 4B**). These findings support a model in which GIGYF2/4EHP-mediated translational repression serves a protective role by limiting the production of potentially toxic protein species upon IR exposure, and that loss of this repression exacerbates cellular damage through the accumulation of aberrant translation products. An exacerbated proteotoxic stress response triggers cell death (62). Therefore, we quantified levels of apoptosis at 48 h post-radiation, the peak of proteotoxic stress response after irradiation based on ATF4 and CHOP expression (**Fig. 4A**), using Annexin/PI staining. While we observed no significant change in basal apoptosis in shCTRL and 4EHP-knockdown cell lines following 2 Gy irradiation, 4EHP depleted cells exhibited a slight but significant increase in early apoptosis indicated by Annexin V-stained cells (3.6±0.4% in shCTR and 6.2±0.9% in sh4EHP, p<0.001) and late-stage apoptosis or secondary necrosis, indicated by Annexin V + PI stained cells (4.1±0.1% in shCTR and 5.2±0.4% in sh4EHP p<0.001; **Fig. 4C & Supp. Fig. 4B)**. Notably, this effect was significantly more pronounced following exposure to 6 Gy irradiation, with 4EHP depletion resulting in a greater difference in annexin V-stained cells compared to control cells (9±2% in shCTR and 21±3% in sh4EHP; p<0.001; **Supp. Fig. 4C & D**).

**Figure 4.**
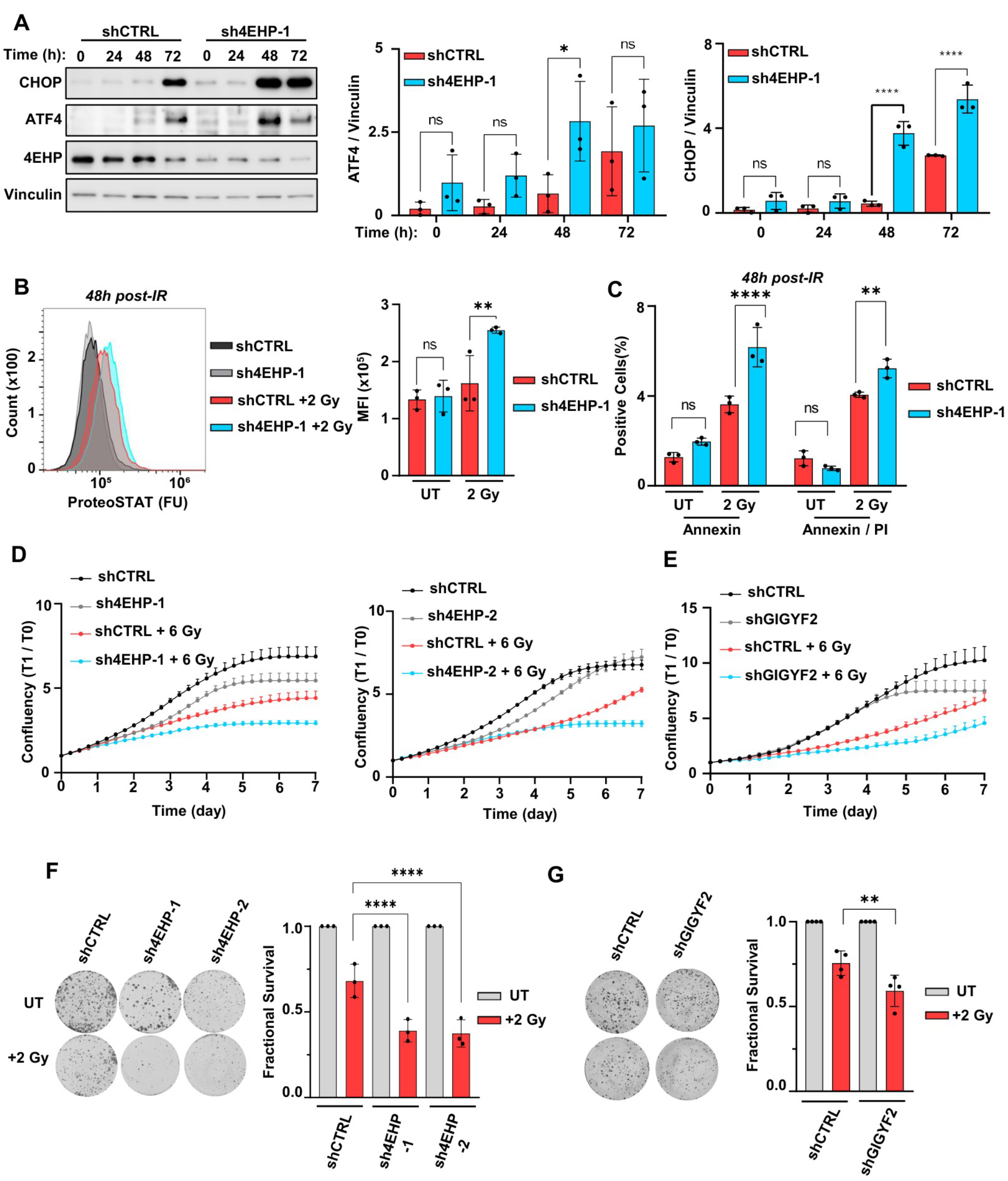
Depletion of 4EHP and GIGYF2 reduces cellular fitness upon IR exposure. (**A**) *Left:* Western blot analysis of expression of the indicated proteins in control (shCTRL) and 4EHP knockdown (sh4EHP-1) U251 cells at the indicated time points following exposure to 2 Gy IR. Vinculin was used as a loading control. *Middle & Right:* Densitometry quantifications of ATF4 and CHOP in the respective conditions. (**B**) *Left:* Analysis of cellular protein aggregation using ProteoSTAT assay. shCTRL and sh4EHP cells were treated with 2 Gy and compared with their untreated counterparts. ProteoSTAT-stained cells were analysed by flow cytometry 48 h after exposure. *Right:* Quantification of the mean fluorescence intensity (MFI) in indicated samples. (**C**) Quantification of % of Annexin V-positive (*Left*) or Annexin V+PI-positive (*Right*) shCTRL and sh4EHP-1 U251 cells 48 h after treatment with 2 Gy IR. (**D & E**) Incucyte SX1 live-cell analysis of proliferation curves derived from % confluency normalised to starting % confluency after 6 Gy irradiation of control (shCTRL) and 4EHP-knockdown (sh4EHP-1 & 2; D) or GIGYF2-knockdown (shGIGYF2) U251 cells (E). Data are presented as representative of three independent replicates. **(F & G)** *Left*: Colony formation assay 10 days after treatment with 2 Gy radiation. *Right*: Quantification of the fractional survival normalised to plating efficiency in shCTRL and 4EHP-knockdown (sh4EHP-1 & 2) U251 cells (F) or shCTRL and GIGYF2-knockdown (shGIGYF2) U251 cells (G). Data are presented as mean ± SD; n=3, one-way Anova; **P<0.01***P<0.001.

Having established that loss of 4EHP disrupted proteostasis and amplified apoptosis in response to IR exposure, we next considered whether cell viability was also affected by depletion of 4EHP- or GIGYF2. We measured the changes in cell growth after IR treatment by Incucyte proliferation assays. Considering the lack of a substantial change in cell growth upon 2 Gy exposure (**Supp. Fig. 1B**) we escalated the IR exposure to 6 Gy, a level sufficient to impose a cell growth defect (**Supp. Fig. 1B**). At this dose, we observed a significantly reduced growth of both 4EHP-KD and GIGYF2-KD U251 cells compared to respective shCTRL cells after IR exposure (33±1%, 17±1%, and 28±2% reductions for sh4EHP-1, sh4EHP-2, and shGIGYF2, respectively; **Fig. 4D & 4E**). Notably, clonogenic survival assays revealed a significant reduction in fractional survival following 2 Gy irradiation in 4EHP- or GIGYF2-depleted cells compared to shCTRL cells (29±5%, 31±5%, 26±4% reduction for sh4EHP-1, sh4EHP-2, and shGIGYF2 respectively; p<0.001; **Fig. 4F & 4G**).

Altogether, these data demonstrate that the GIGYF2/4EHP translational repressor complex is critical for maintenance of cellular homeostasis and viability through reducing the proteotoxic burden in response to IR exposure.

### Ionizing radiation augments the interaction between RBPs and GIGYF2/4EHP complex

We next aimed to characterize the mechanism by which the GIGYF2/4EHP translational repression is employed after IR treatment. 4EHP is a relatively weak cap-binding protein (30-100 fold lower than eIF4E) (34). To achieve a competitive advantage over the canonical cap-binding translation factor eIF4E, 4EHP must be recruited to its target mRNAs by RBPs such as ZFP36/TTP, ZNF598 and TNRC6 proteins (components of miRISC) via GIGYF2 (24) (**Fig. 3A**). We therefore hypothesised that recruitment of GIGYF2/4EHP complex to its target mRNAs upon IR treatment is mediated by one or both of the following mechanisms: i. changes in the binding activity of the GIGYF2/4EHP-interacting RBPs to target mRNAs; ii. Changes in the protein-protein interactions between GIGYF2/4EHP and RBPs (**Fig. 5A**).

**Figure 5.**
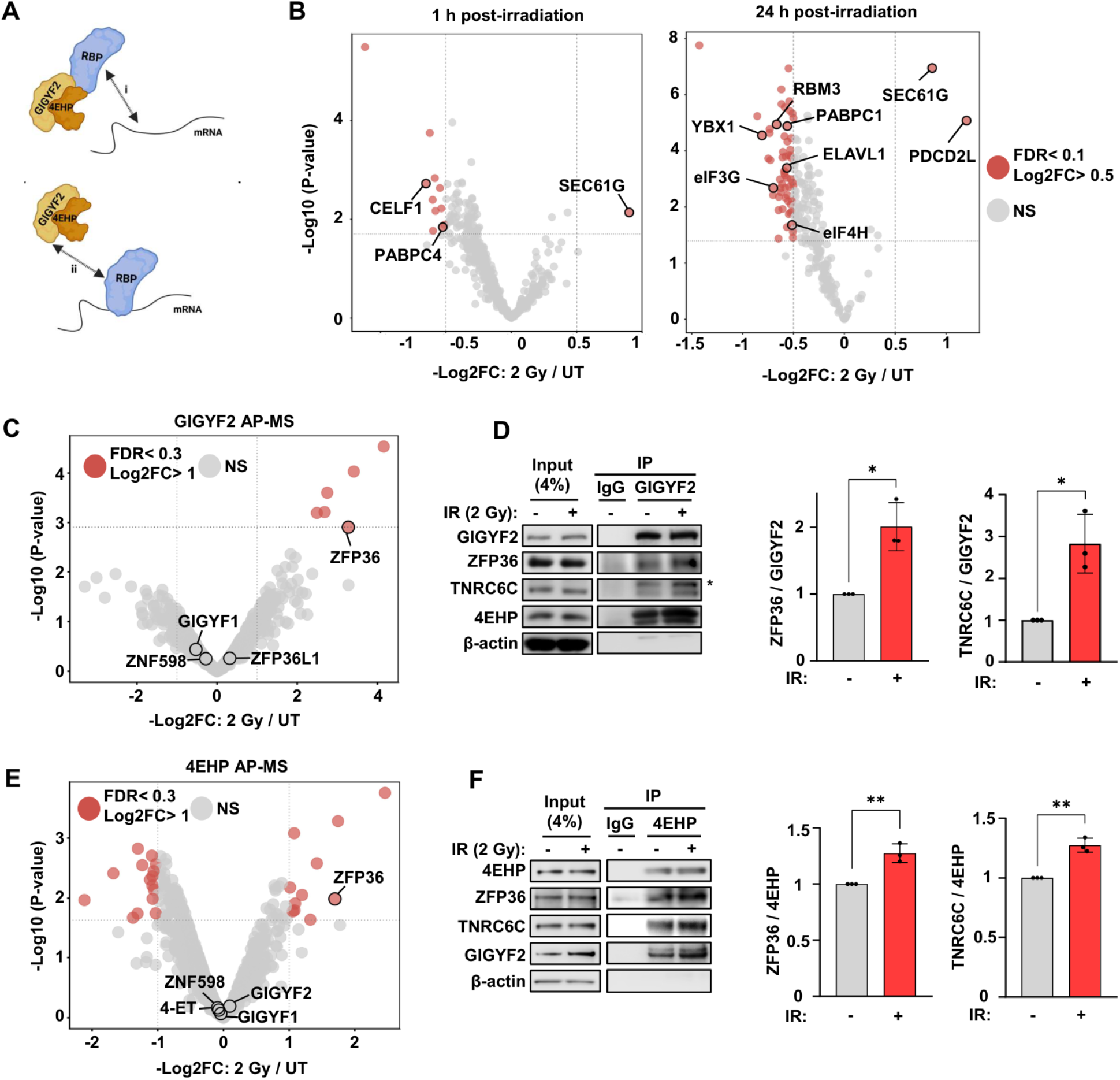
IR rapidly enhances the interactions between GIGYF2/4EHP and partner RBPs. (**A**) Schematic model of two alternative proposed mechanisms responsible for the enhanced translational repression by the GIGYF2/4EHP complex upon IR exposure: i. changes in the interactions between mRNAs and RBPs, ii. Changes in the interactions between GIGYF2/4EHP and RBPs. (**B**) Comparative RNA interactome capture analysis of poly(A) RNA-bound proteins in U251 cells at 1 h (*Left*) or 24 h (*Right*) post-treatment with 2 Gy IR. Volcano plots show the Log2 fold change and the -Log10 p-value for UV-enriched proteins 1 and 24 h following 2 Gy irradiation, relative to time-matched untreated controls. n=6 independent replicates. Proteins with Log2FC>0.5 and FDR<0.1 are coloured red and proteins with known roles in translational regulation and radiation response are annotated. (**C**) Volcano plots showing the Log2 fold-change and the -Log10 p-value of proteins enriched by GIGYF2 immunoprecipitation 1 h following 2 Gy irradiation, relative to time-matched untreated controls. Proteins with Log2FC>1 and FDR<0.3 are coloured in red. Known direct interactors of GIGYF2 are annotated. n=3 independent replicates (**D**) *Left:* Co-IP for detection of interaction between endogenous GIGYF2 and the indicated proteins in U251 cells 1 h post-treatment with 2 Gy IR or time-matched untreated control. *Right:* Densitometry quantifications of ZFP36 and TNRC6C normalized to GIGYF2 pulldown in the respective cells presented as mean ± SD; n=3, one-way Anova; *p<0.05. Astrix denotes non-specific band. (**E**) Volcano plots showing the Log2 fold-change and the -Log10 p-value for proteins enriched by 4EHP immunoprecipitation 1 h following 2 Gy irradiation, relative to time-matched untreated controls. Proteins with Log2FC>1 and FDR<0.3 are coloured in red, and known direct interactors of GIGYF2 are annotated. n=3 independent replicates. (**F**) *Left:* Co-IP for detection of interaction between endogenous 4EHP and the indicated proteins in U251 cells 1 h following 2 Gy irradiation or time-matched untreated control. *Right:* Densitometry quantifications of ZFP36 and TNRC6C normalized to 4EHP pulldown in the respective cells presented as mean ± SD; n=3 independent replicates, one-way Anova; **p<0.01.

To test whether GIGYF2/4EHP recruitment is driven by changes in the interactions between relevant RBPs and mRNAs, we assessed the net mRNA association of all RBPs at 1 and 24 hour post 2 Gy IR treatment compared with untreated cells via comparative RNA-Interactome Capture (cRIC) assay (63). In brief, RIC uses UV to promote *in situ*, zero distance protein-RNA crosslinks between proteins and RNA, followed by denaturing purification of polyadenylated RNA (and covalently bound proteins) with oligo(dT) and quantitative proteomics. We identified 452 proteins enriched upon UV cross-linking across all experimental conditions (Log2FC>0.5, FDR<0.05; **Supp. Fig. 5A & Supp. Table 1**), ∼95% of which have been previously annotated as RBPs in ≥3 of studies collated in RBPbase (64) (**Supp. Fig. 5B**). This validates the high degree of specificity of the assay in detecting mRNA-bound RBPs (RBPome). After successfully defining the RBPome in untreated and irradiated U251 cells by UV enrichment, we sought to assess the changes in the RBPome after irradiation. the 79 RBPs changed upon radiation, only 3 RBPs (ACTG1, SEC61G, and PDCD2L) exhibited increased mRNA binding, with only SEC61G displaying consistent upregulation at 1 and 24 h after IR treatment (Log2FC>0.5, FDR<0.1; **Fig. 5B & Supp. Table 2**). The majority of RBPs displayed reduced mRNA binding upon irradiation, where we detected 15 and 69 RBPs with reduced mRNA binding at 1 and 24 h post-IR treatment, respectively (**Fig. 5B & Supp. Table 2**). These included several RBPs with known roles in regulation of mRNA translation and the response to radiation such as eIF3G (65,66), eIF4H (67), CELF1 (68), PABPC4 (69), PABPC1 (69), RBM3 (70), YBX1 (71), and HuR/ELAVL1 (72). Notably, HuR/ELAVL1 has been previously shown to dissociate from mRNA following IR exposure to confer radioprotection in colorectal cancer cells (72). PABPC1/4 have been shown to localize to the nucleus, sequestering from the bulk of mRNAs, following UV irradiation (73,74). Together, these data are consistent with the observed rapid and persistent translational repression of mRNA translation in GBM cells following IR treatment (**Fig. 1**) and further support the utility of the cRIC approach for identifying changes in RBPome in response to IR. However, we were unable to identify differential mRNA binding for any known interactors of the GIGYF2/4EHP complex at 1 hour or 24 hours post-IR. This suggests that GIGYF2/4EHP recruitment to and translational repression of their target mRNAs upon IR treatment cannot be explained by changes in the interactions between the relevant RBPs and mRNAs.

Therefore, we assessed if the recruitment of GIGYF2/4EHP upon radiation is mediated by changes in their interaction with RBPs. We performed an Affinity Purified-Mass-Spec (AP-MS) assay by pulldown of endogenous GIGYF2 protein to identify the changes in its protein-protein interactome 1 h post-radiation (2 Gy), compared with untreated cells. Immunoprecipitation was performed in presence of ribonuclease to remove false-positives due to potential RNA-bridged interactions. Significant enrichment of several known interactors of GIGYF2, including ZNF598 (27,31), and ZFP36L1 (75,76) (Log2FC>1, FDR<0.05; **Supp. Fig 5C & Supp. Table 3**), in both untreated and irradiated cells highlighted the robustness of our AP-MS assay in identifying the GIGYF2 interacting proteins. As expected, Gene Ontology (GO) analysis of the GIGYF2 interactome revealed significant enrichment for translation-related biological processes (**Supp. Fig. 5D & Supp. Table 4**). When comparing the changes in GIGYF2 interactome between the untreated and IR-treated cells, among the most prominent changes was a significantly increased interaction with ZFP36/TTP (Log2FC=3.3; **Fig 5C & Supp. Table 3**), a known recruiter of the GIGYF2/4EHP complex (28). Notably, this interaction was only detected following irradiation (**Supp. Fig 5C**). We further validated changes in known direct interactors of GIGYF2 by western blot following Co-IP. Consistent with our AP-MS data, IR treatment led to a significantly increased interaction between GIGYF2 and ZFP36 (2±0.3 fold increase, p<0.05; **Fig. 5D**). Our Co-IP/western blot additionally revealed significantly increased interaction between GIGYF2 and TNRC6C after radiation (2.8±0.7 fold increase, p<0.05; **Fig. 5D**). We complemented these findings via AP-MS for endogenous 4EHP in U251 cell 1 hour post-radiation, compared with the untreated cells. We identified several known 4EHP interactors including GIGYF1, GIGYF2, ZNF598 and 4-ET among the preys identified in both untreated and irradiated cells (Log2FC>1, FDR<0.05; **Supp Fig. 5E & Supp. Table 5**). When comparing the effect of radiation on 4EHP interactome, similar to the GIGFYF2 interactome (**Fig. 5C**), we noted an increase in 4EHP-ZFP36 interaction (Log2FC= 1.7; **Fig. 5E & Supp. Table 5**). This was further validated by western blotting following 4EHP pulldown where we observed a 30% increased 4EHP-ZFP36 interaction following irradiation (p<0.05; **Fig. 5F**). Similarly, we detected a significant increase in the interaction between 4EHP and TNRC6C upon IR exposure (3.6±0.6 fold increase, p<0.05; **Fig. 5F**). Consistent with a GIGYF2 bridged interaction between 4EHP and RBPs such as TNRC6 or ZFP36 (**Fig. 3A**) (32,77), we observed a lower level of enrichment of TNRC6C or ZFP36 proteins in 4EHP pull-downs (**Fig. 5F**) compared with GIGYF2 pull-downs after irradiation (**Fig. 5D**) by western blotting. In support of this, we observed a reduced enrichment of ZFP36 after radiation from AP-MS of 4EHP compared to GIGYF2 (Log2FC:3.3 vs.7 for 4EHP and GIGYF2 pulldown, respectively: **Fig 5C & D)**.

Collectively, these data indicate that upon IR exposure increased protein-protein interactions with RBPs facilitates the recruitment of the GIGYF2/4EHP complex to mediate translational repression without widespread shifts in the binding of these RBPs to mRNA targets (**Fig. 6**).

**Figure 6.**
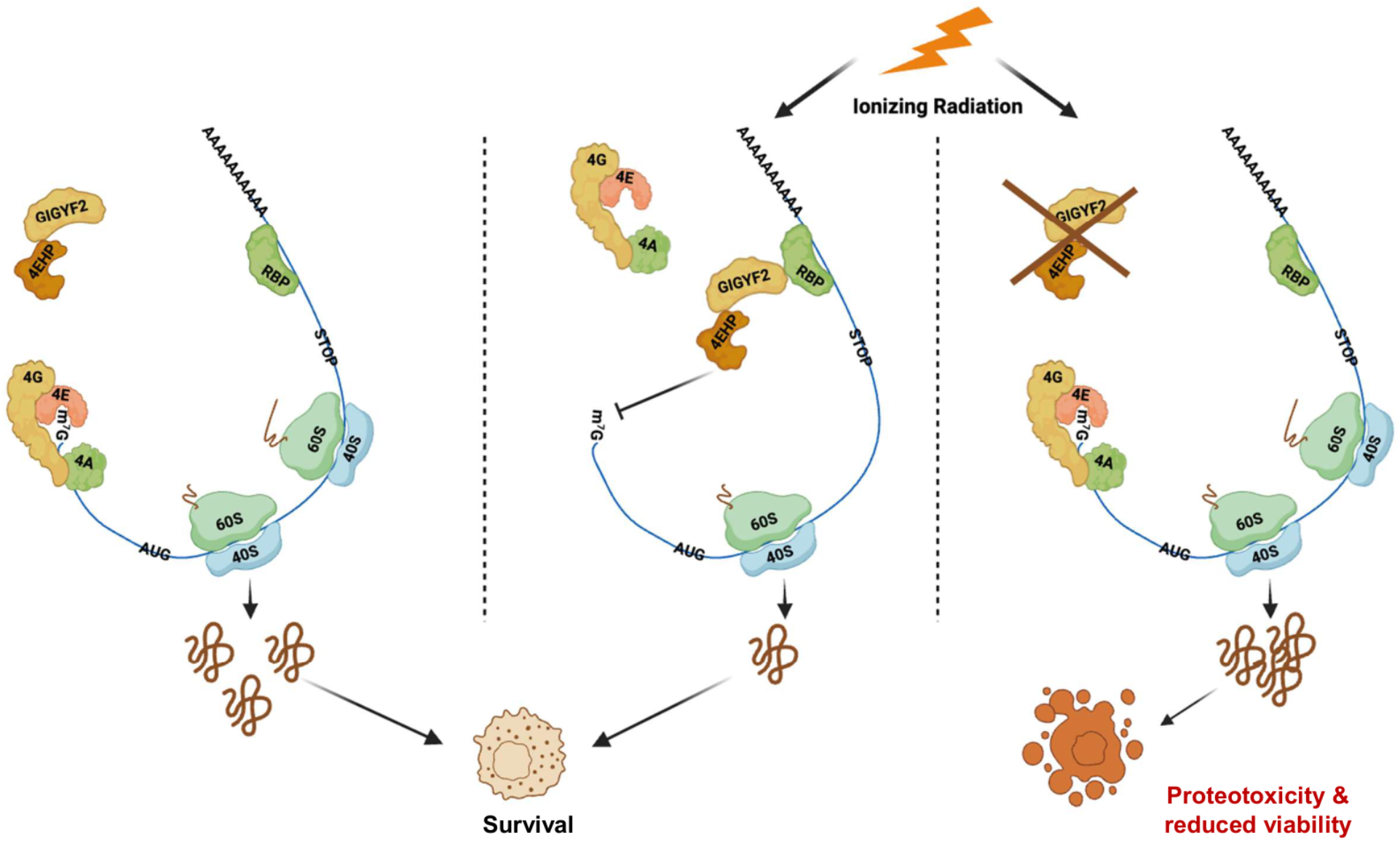
GIGYF2/4EHP-mediated translational repression maintains cell survival upon ionizing radiation. Proposed model for the mechanism of translational repression and maintenance of cellular homeostasis following IR exposure, mediated by the GIGYF2/4EHP complex. *Left*: When the 4EHP-GIGYF2 complex is not recruited to its targeted transcripts eIF4E, along with eIF4G and eIF4A, binds to the cap and initiate translation. *Middle:* Upon IR exposure, the GIGYF2/4EHP translational repressor complex is recruited to specific target mRNAs through enhanced binding to RNA-binding proteins (RBPs), such as ZFP36. This recruitment reduces the overall translational load on these mRNAs and helps prevent aberrant translation events that would otherwise lead to proteotoxicity. By limiting the production of defective or surplus proteins, the complex supports cellular homeostasis and protects against IR-induced stress. *Right*: In the absence of the GIGYF2/4EHP complex, the rapid translational repression triggered by IR is abrogated. Consequently, translation continues unchecked, leading to the accumulation of abnormal translation products. These products overwhelm the proteostasis machinery, resulting in increased proteotoxicity and a failure to maintain cellular homeostasis upon IR exposure.

## Discussion

Regulation of mRNA translation is a critical component of the cellular stress response. Here, we describe a mechanism of cellular response to ionizing radiation that operates through rapid and coordinated modulation of a substantial fraction of cellular mRNA translation output to maintain cellular homeostasis under stress conditions. We demonstrate that glioblastoma cells rapidly repress mRNA translation in response to a clinically relevant dose of ionizing radiation to maintain cellular homeostasis. The rapid response is largely mediated by engagement of a translational repressor complex composed of the mRNA cap-binding protein 4EHP and its binding partner GIGYF2. This translational repression protects cells from radiation-induced toxicity by limiting proteotoxic stress. Abrogation of this mechanism leads to higher susceptibility of the glioblastoma cells to radiation-induced proteotoxicity, thus compromising cell viability.

We demonstrate that recruitment of the GIGYF2/4EHP complex to target mRNAs is facilitated by enhanced interactions with RBPs (e.g. ZFP36 and TNRC6C) that are already associated with these transcripts as evidenced by the increased binding of these proteins to GIGYF2 and 4EHP but lack of change in their mRNA binding upon radiation (**Fig. 5**). This mechanism allows rapid reprogramming of mRNA translation by leveraging pre-bound RBPs, rather than relying on slower processes such as changes in gene expression or widespread remodeling of RBP-mRNA interactions. GIGYF2 directly interacts with several key RBPs, including miRISC components (e.g. TNRC6C) and ZFP36 (tristetraprolin or TTP), which regulate target mRNAs with varying efficiencies (78,79). Regulated recruitment of the GIGYF2-4EHP complex to these RBPs therefore enables a rapid, scalable, and finely tuned program of transcript-specific translational repression.

Although 4EHP directly interacts with the mRNA 5’ cap, its affinity for the cap is much weaker than that of its more potent cap-binding homologue, eIF4E. Consequently, recruitment of 4EHP to target mRNAs relies on initial interactions with RBPs, such as ZFP36, ZNF598, and the miRISC complex, that bind to the 3’ UTR or coding sequence (CDS) of the same mRNA. However, 4EHP does not typically bind directly to these RBPs. Instead, previous studies have identified three intermediary proteins - GIGYF1, GIGYF2, and 4E-T - that bridge 4EHP to RBPs (25,31,80). The binding of these proteins to 4EHP is competitive, thus creating a range of possibilities through which RBPs can recruit 4EHP to their target mRNAs via any of the three intermediates. Once recruited to target mRNAs, through a combination of increased local concentration and enhanced affinity for the 5’ cap (25), 4EHP displaces eIF4E and thereby blocks translation initiation. Crucially, given the diverse array of RBPs that recruit 4EHP via GIGYF2, GIGYF1, or 4E-T as intermediaries, the precise set of mRNAs affected by this mechanism likely differs across cell types and even within the same cell under different contexts (e.g., distinct cell cycle stages). This offers the cell considerable flexibility to selectively reduce the translation of specific mRNAs to varying degrees, thereby maintaining cellular homeostasis in response to the radiation-induced stress. In this study, we specifically assessed the role of the GIGYF2/4EHP complex in the cellular response to ionizing radiation. However, it remains plausible that the GIGYF1/4EHP and 4E-T/4EHP complexes may also play similar or parallel roles under these conditions, a possibility that should be addressed in future studies.

The recruitment of GIGYF2 to specific mRNAs is mediated through interactions with partner RBPs. The best-characterized mode of such interaction involves the GYF (glycine-tyrosine-phenylalanine) domain, a versatile adaptor module that recognizes proline-rich sequences (PRS) in partner proteins. Notably, only three GYF domain-containing proteins are encoded in the human genome: CD2BP2, GIGYF1, and GIGYF2 (81). PRS motifs are found in several RBPs, including ZFP36, ZNF598, and the TNRC6 (GW182) family of proteins, which are central to post-transcriptional gene regulation. The specificity of 4EHP-mediated translational repression is thought to be conferred by the GYF domain of GIGYF1/2 proteins, which acts as an adaptor bridging PRS-containing RBPs to the 4EHP translational repressor. Importantly, interactions between GIGYF1/2 and their RBP partners can be dynamically regulated via phosphorylation of residues flanking the GYF domain. For instance, a previous study demonstrated that p38 kinase-mediated phosphorylation of Ser638 on GIGYF1 upon lipopolysaccharide (LPS) treatment modulates its interactions with ZFP36 and ZNF598 (82). Although a comparable modification on GIGYF2 was not detected in that study, the high degree of sequence conservation in this region suggests that similar post-translational events may also regulate GYF:PRS-mediated binding of GIGYF2 to its partner RBPs. More recent work revealed an inverse correlation between phosphorylation of GIGYF2 in its N-terminal region, near the 4EHP-binding motif, and its recruitment to mRNAs in interferon-treated cells (61). Collectively, these findings support a model in which post-translational modifications govern the binding of GIGYF1/2 to partner RBPs and the subsequent recruitment of these RBPs to target mRNAs in a stress- and context-dependent manner. Given the crucial role of the kinases in orchestrating the response to radiation (83), it is plausible that analogous post-translational mechanisms contribute to the rapid, IR-induced increase in interactions between the GIGYF2/4EHP complex and its cognate RBPs, thereby driving translational repression. Future studies aimed at mapping IR-induced phosphorylation events on GIGYF2 and its interacting partners will be essential to fully elucidate the regulatory logic underlying this stress response pathway.

While this study primarily focused on well-established direct interactors of the GIGYF2/4EHP complex, several other proteins also exhibit increased mRNA binding or increased binding to GIGYF2/4EHP after irradiation. We cannot exclude the possibility that some of these proteins may also bind to and recruit the GIGYF2/4EHP complex to mRNAs, thereby contributing to GIGYF2/4EHP-mediated translational repression following IR treatment. For example, we observed increased mRNA binding of SEC61G, the small gamma subunit of the Sec61 translocon complex, which serves as the central channel for protein translocation into the endoplasmic reticulum (ER), at both 1 hour and 24 hours post-IR (**Fig. 5B**). Although no direct interaction between SEC61G and the GIGYF2/4EHP complex has been reported to date, a recent study demonstrated that Sec61β, another ER translocon subunit, inhibits cap-dependent translation of ER stress-induced pre-emptive quality control (ERpQC) substrates by recruiting 4EHP via the E3 ubiquitin ligase ARIH1 (37). Despite observing increased Sec61 binding to mRNA, we found no interaction between 4EHP and ARIH1 and while we detected an interaction between 4EHP and Sec61β, this interaction was stable upon irradiation. Collectively, this suggests that 4EHP’s interaction with Sec61β is unlikely to drive its early (1 h) recruitment to mRNAs to mediate translational repression. However, we cannot rule out that modulation of this interaction at later (>1 h) timepoints not tested here may contribute to 4EHP-dependent attenuation of proteotoxic stress observed at 48 hours (**Fig. 4**).

We demonstrate that GIGYF2/4EHP-mediated early translational repression may be important for averting activation of the ER stress response following irradiation, as evidenced by the exacerbated formation of protein aggregates and increased expression of ATF4 and CHOP in 4EHP-depleted cells with compromised repression of translation post-IR. However, it is plausible that this mechanism is only partially responsible for the reduced viability of 4EHP- or GIGYF2-depleted cells upon IR exposure. Indeed, the GIGYF2/4EHP complex has been reported to regulate several other critical processes, including the repression of translation of microRNA target mRNAs, mRNAs with collided ribosomes, mRNAs subjected to nonsense-mediated decay, non-optimal mRNAs, and those targeted by the ZFP36 family of RNA-binding proteins (25,26,30,32,36,60,84–86). Moreover, the complex plays an essential role in repressing the translation of key factors involved in the activation of the innate immune response, such as cGAS and proinflammatory cytokines (e.g., interferon-β) (35,36,77). Therefore, other cytotoxic mechanisms beyond ER proteotoxicity may also contribute to the reduced viability of GIGYF2/4EHP-deficient cells following IR exposure.

Importantly, the present study was conducted exclusively using glioblastoma cell models. Given the extensive array of mechanisms influenced by the GIGYF2/4EHP complex, it is possible that other cell types -particularly those with different baseline stress responses or immune activity-may exhibit additional or distinct phenotypes. A more comprehensive characterization of changes in cellular mRNA translation and associated pathways, will likely reveal the broader role of this complex in regulating both cellular and organismal responses to ionizing radiation.

While our analysis of general mRNA translation and new protein synthesis revealed that 4EHP or GIGYF2 knockdown has a sizeable effect on translation after irradiation, it still only partially restores the reduced mRNA translation and protein synthesis 1 hour after irradiation (**Fig. 3**). This incomplete recovery may be explained by the residual 4EHP or GIGYF2 proteins in knockdown cells, which could continue to exert partial translational repression. However, we cannot rule out the contribution of additional, 4EHP- or GIGYF2-independent pathways to this phenomenon. Supporting this possibility, we observed significantly reduced mRNA binding of several translation-promoting RBPs including PABPC4 (87), CELF1 (68), and PYM1 (88), 1 hour following irradiation, that may plausibly contribute to GIGYF2-4EHP-indendependent repression (**Fig. 5B**). Notably, a broader and more pronounced downregulation of mRNA binding was observed at 24 hours post-treatment compared to 1 hour, suggesting a temporal shift in RBP-mRNA interaction dynamics. These changes may arise from altered expression profiles of either the mRNAs or the RBPs themselves, or from the activation of distinct translational regulatory mechanisms at different stages following IR. The functional roles of the IR-perturbed RBPs identified here in coordinating the post-transcriptional response to IR remains an important question for future investigation, although it is feasible that their reduced binding may contribute to/or be a product of a sustained block on translation after irradiation (**Fig 1A**).

Our study has demonstrated RBP-directed recruitment of the GIGYF2-4EHP complex following irradiation, however the precise upstream lesions, whether directly on the RNA or mediated through cellular signaling cascades triggered by DNA damage, remain to be resolved. Future studies employing targeted approaches to uncouple these processes will be necessary to delineate the primary triggers driving GIGYF2/4EHP-mediated translational control in response to radiation.

Collectively, our findings position translational repression not merely as a downstream consequence of ionizing radiation IR-induced cellular stress, but as a critical and functionally significant arm of the cellular response that actively shapes cell fate following genotoxic injury. We further demonstrate the critical role of the GIGYF2/4EHP translational repressor complex in orchestrating this adaptive response. By attenuating selectively regulating specific mRNA targets, this complex helps cells manage proteotoxic stress, limit the production of aberrant translation products, and ultimately influence cell viability following irradiation. Although our study was primarily focused on elucidating the fundamental biological pathways governing the early cellular response to IR, these findings may carry potentially significant implications for clinically relevant contexts such as radiation therapy. For instance, modulating the activity of the GIGYF2/4EHP axis could represent a novel strategy to sensitize tumor cells to irradiation by preventing them from mounting an effective protective translational response. Alternatively, baseline expression levels or radiation-induced activation status of this complex may serve as predictive biomarkers for responses to radiotherapy. Future studies exploring the therapeutic potential of targeting the GIGYF2/4EHP pathway, as well as its role in the cellular response to other genotoxic stresses, will be important next steps.

## Data Availability

All data are incorporated into the article and its online supplementary material.

## Author contributions

**Conceptualization:** T.M. and S.M.J.

**Investigation:** T.M., D.M., H.A., O.O., P.H.S., S.C., A.K., Y.C.

**Methodology:** A.C., N.S., R.G.

**Resources:** R.L., H.A., P.N., K.B.

**Data curation & analysis:** T.M., T.S., C.A.

**Validation:** S.M.

**Writing - original draft:** T.M. and S.M.J.

**Writing - review & editing:** T.M., D.M., O.O., P.H.S., S.C., A.K., Y.C., T.S., R.L., H.A., P.N., C.A., K.B., S.M., A.C., N.S., R.G. S.M.J.

**Project administration:** T.M. and S.M.J.

**Funding acquisition:** S.M.J.

**Supervision:** S.M.J.

## Acknowledgment

We thank Dr. Mihaela Pettigrew for enabling access to the Multirad160 device.

## Supplementary Figure Legends

**Supplementary Figure 1: Ionizing radiation represses mRNA translation in GBM cells**. (**A**) *Left:* Western blot analysis of expression of cleaved PARP (c-PARP) using lysates derived from U251 cells 24 h and 48 h after 2 Gy irradiation. Vinculin was used as a loading control. *Right:* Densitometric quantitation of the cleaved-PARP/total-PARP signals normalized to Vinculin for each replicate and presented as mean ± SD; n=3; **p<0.01, one-way Anova. (**B**) Incucyte SX1 live-cell analysis of proliferation curves derived from % confluency normalised to starting % confluency after 2 Gy or 6 Gy irradiation of U251 cells. Data are presented as mean ± SD; n=3, two-way Anova; ns=non-significant, ***p<0.001. (**C-H**) Representative polysome profiling traces from U251 (C-F) and U87 cells (G & H) at the indicated time points following 2 Gy irradiation. (**I-K**) Quantification of nascent protein synthesis measured by metabolic pulse labelling with the methionine analogue L-homopropargylglycine (HPG) 1 h after irradiation with in 0.1 Gy (I), 1 Gy (J), and 6 Gy (K). Normalized fluorescence intensity was calculated by deduction of the unstained fluorescence followed by division by untreated fluorescence. Data are presented as mean ± SD; n=3, two-way Anova; ns=non-significant, *p<0.05, ***p<0.001. (**L**) Representative DNA content histograms of U251 cells analyzed by PI staining in U251 cells 1 h following 2 Gy irradiation. (**M**) Quantification of cell cycle distribution in (I). Data are presented as mean ± SD; n=3, one-way Anova; ns=non-significant.

**Supplementary Figure 2: mTOR and ISR-independent cellular response to IR in GBM cells**. (**A & B**) *Top:* Western blot analysis of phosphorylation of RPS6 (Serine 240/244) in U251 (A) and U87 cells (B) following 2 Gy irradiation. Torin-1 (15 nM) treatment for 24 h was used as a control. Vinculin was used as a loading control. *Bottom:* Densitometry quantifications of p-RPS6/RPS6 in the respective cells. (**C**) Quantification of new protein synthesis by Surface Sensing of Translation (SUnSET) assay in U251 cells pre-treated with ISRIB or DMSO for 1 h prior to tunicamycin treatment with indicated dosages for 4 h. CHX treatment was used as a control. Cells were collected following a 15 min pulse with 2 μg/mL puromycin. Vinculin was used as a loading control. (**D**) Colony formation assay with U251 cells. Cells were pre-treated with the indicated dose of ISRIB for 1 h followed by 2 Gy irradiation. 10 days post-seeding cells were fixed and fractional survival normalized to plating efficiency. Data are presented as mean ± SD; n=3, one-way Anova; ns=non-significant, *p<0.05.

**Supplementary Figure 3. Depletion of 4EHP and GIGYF2 reverses the translational repression upon IR exposure**. (**A**) Western blot analysis of expression of 4EHP in U251 cells with stable expression of two independent shRNAs against 4EHP (sh4EHP-1 and sh4EHP-2) or a control shRNA (shCTRL). (**B**) Western blot analysis of expression of GIGYF2 in U251 cells with stable expression of a shRNA against GIGYF2 (shGIGYF2) or shCTRL. (**C**) Western blot analysis of expression of 4EHP in U87 cells with stable expression of sh4EHP-1 or shCTRL. (**D**) Quantification of nascent protein synthesis in shCTRL and sh4EHP U87 cells measured by metabolic pulse labelling with the methionine analogue L-homopropargylglycine (HPG), 1 h post-treatment with 2 Gy radiation. Signals were normalized against the unstained samples for each group. Data are presented as mean ± SD; n=3, one-way Anova; ns=non-significant, *p<0.05.

**Supplementary Figure 4. 4EHP contributes to the maintenance of cellular fitness upon IR exposure**. (**A**) Analysis of cellular protein aggregation using ProteoSTAT assay, cells were analysed by flow cytometry 16 h after treatment with 5 µM of the proteasome inhibitor MG132. (**B**) Representative flow cytometry density plots from Annexin V/PI stained shCTRL and sh4EHP-1 U251 cells 48 h after treatment with 2 Gy IR. (**C**) Representative flow cytometry density plots from Annexin V/PI stained shCTRL and sh4EHP-1 U251 cells 48 h after treatment with 6 Gy IR. (**D**) Quantification of % of Annexin V-positive (*Left*) or Annexin V+PI-positive (*Right*) shCTRL and sh4EHP-1 U251 cells 48 h after treatment with 6 Gy IR.

**Supplementary Figure 5. IR-induced changes in RBPome and GIGYF2 or 4EHP protein interactomes in U251 cells**. (**A**) Volcano plot showing the Log2 fold-change and the p-value for protein enrichment by UV crosslinking in the untreated U251 cells. n= 6 independent replicates. Proteins with FDR<0.05 and Log2FC>0.5 are coloured red. (**B**) Bar chart of the proportion of UV-enriched proteins (FDR<0. and Log2FC>0.5) across conditions that were identified as RBPs in ≥3 studies collated in RBPbase database. (**C**) Volcano plots showing the Log2 fold-change and the p-value for proteins enrichment by endogenous GIGYF2 immunoprecipitation over IgG control in untreated U251 cells (*Left*), or 1 h after 2 Gy irradiation (*Right*). Proteins with FDR<0.05 are coloured in red. n=3 independent replicates. (**D**) Gene ontology analysis of biological pathways related to proteins significantly enriched (FDR<0.05) by GIGYF2 immunoprecipitation over IgG controls. (**E**) Volcano plots showing the Log2 fold-change and the p-value for proteins enrichment by endogenous 4EHP immunoprecipitation over IgG control in untreated U251 cells (*Left*), or 1 h after 2 Gy irradiation (*Right*). Proteins with FDR<0.05 are coloured in red. n=3 independent replicates.

**Supplementary Figures 6-12:** Uncropped images of western blots used in this manuscript.

## Supplementary Tables

**Supplementary Tables 1: UV-enriched proteins from RNA-Interactome Capture (RIC)**. Proteins significantly enriched in UV-crosslinked U251 cells relative to non-crosslinked control cells, as determined by RNA-interactome capture coupled to quantitative mass spectrometry. Differential enrichment analysis was performed using the limma R. package to produce four tables for UV-enrichment under each condition. The tables report the log2 fold change (logFC), average log2 expression intensity (AveExpr), moderated t-statistic (t), nominal p-value (P.Value), Benjamini–Hochberg adjusted p-value (adj.P.Val), log-odds of differential enrichment (B), and gene symbol for each identified protein. Positive logFC values indicate proteins enriched in UV-crosslinked samples and therefore associated with RNA in U251 cells. The last table reports all proteins significantly enriched by UV crosslinking (Log2FC>0.5, FDR<0.1) under any of the experimental conditions, which was used as a list to filter for RNA-binding proteins for further analysis of radiation treatment effect.

**Supplementary Tables 2: Radiation treatment effect from comparative RNA-interactome capture**. Proteins significantly enriched in irradiated UV-crosslinked U251 cells relative to untreated UV-crosslinked control cells, as determined by comparative RNA-interactome capture coupled to quantitative mass spectrometry. Differential enrichment analysis was performed using the limma R. package to produce two tables for the effect of radiation on the population of UV-crosslinked proteins. The tables report the log2 fold change (logFC), average log2 expression intensity (AveExpr), moderated t-statistic (t), nominal p-value (P.Value), Benjamini–Hochberg adjusted p-value (adj.P.Val), log-odds of differential enrichment (B), and gene symbol for each identified protein. The penultimate column indicates whether the protein was defined as significantly enriched (Log2FC>0.5, FDR<0.1) by UV irradiation from the analysis described in Supplementary Table 1. The final column indicates whether the protein has been previously annotated as an RNA-binding protein in ≥3 studies collated in RBPbase database. Positive logFC values indicate proteins enriched in irradiated UV-crosslinked samples.

**Supplementary Tables 3: GIGYF2 AP-MS after radiation**. Proteins significantly enriched in endogenous GIGYF2 co-immunoprecipitation relative to IgG controls, as determined by quantitative mass spectrometry. Differential enrichment analysis was performed using the limma R. package to produce two tables for protein enrichment over IgG controls. The final table reports the differential enrichment analysis for irradiated (2 Gy, 1 h) GIGYF2 co-IP compared to untreated co-IP, following filtering for any interacting proteins significantly enriched over IgG. The tables report the log2 fold change (logFC), average log2 expression intensity (AveExpr), moderated t-statistic (t), nominal p-value (P.Value), Benjamini–Hochberg adjusted p-value (adj.P.Val), log-odds of differential enrichment (B), and gene symbol for each identified protein.

**Supplementary Tables 4: GO analysis of GIGYF2 AP-MS interactors**. Gene Ontology (GO) enrichment analysis performed on proteins significantly enriched in GIGYF2 affinity purification–mass spectrometry (AP–MS) experiments compared to IgG control immunoprecipitations. Enrichment analysis was performed using clusterProfiler, and overrepresentation was assessed against the human proteome background. Each row corresponds to an enriched GO term and includes the number of input proteins associated with each term (Count/GeneRatio), the background ratio (BgRatio), enrichment statistics (RichFactor, Fold Enrichment, and z-score), and significance metrics (p-value, adjusted p-value, and q-value). The “geneID” column lists the proteins contributing to each enriched term.

**Supplementary Tables 5: 4EHP AP-MS after radiation**. Proteins significantly enriched in endogenous 4EHP co-immunoprecipitation relative to IgG controls, as determined by quantitative mass spectrometry. Differential enrichment analysis was performed using the limma R. package to produce two tables for protein enrichment over IgG controls. The final table reports the differential enrichment analysis for irradiated (2 Gy, 1 h) 4EHP co-IP compared to untreated co-IP, following filtering for any interacting proteins significantly enriched over IgG. The tables report the log2 fold change (logFC), average log2 expression intensity (AveExpr), moderated t-statistic (t), nominal p-value (P.Value), Benjamini–Hochberg adjusted p-value (adj.P.Val), log-odds of differential enrichment (B), and gene symbol for each identified protein.

