## Supplementary Figures 1-12 for "GIGYF2/4EHP-Mediated Translational Attenuation Maintains Cellular Homeostasis Following Ionizing Radiation"

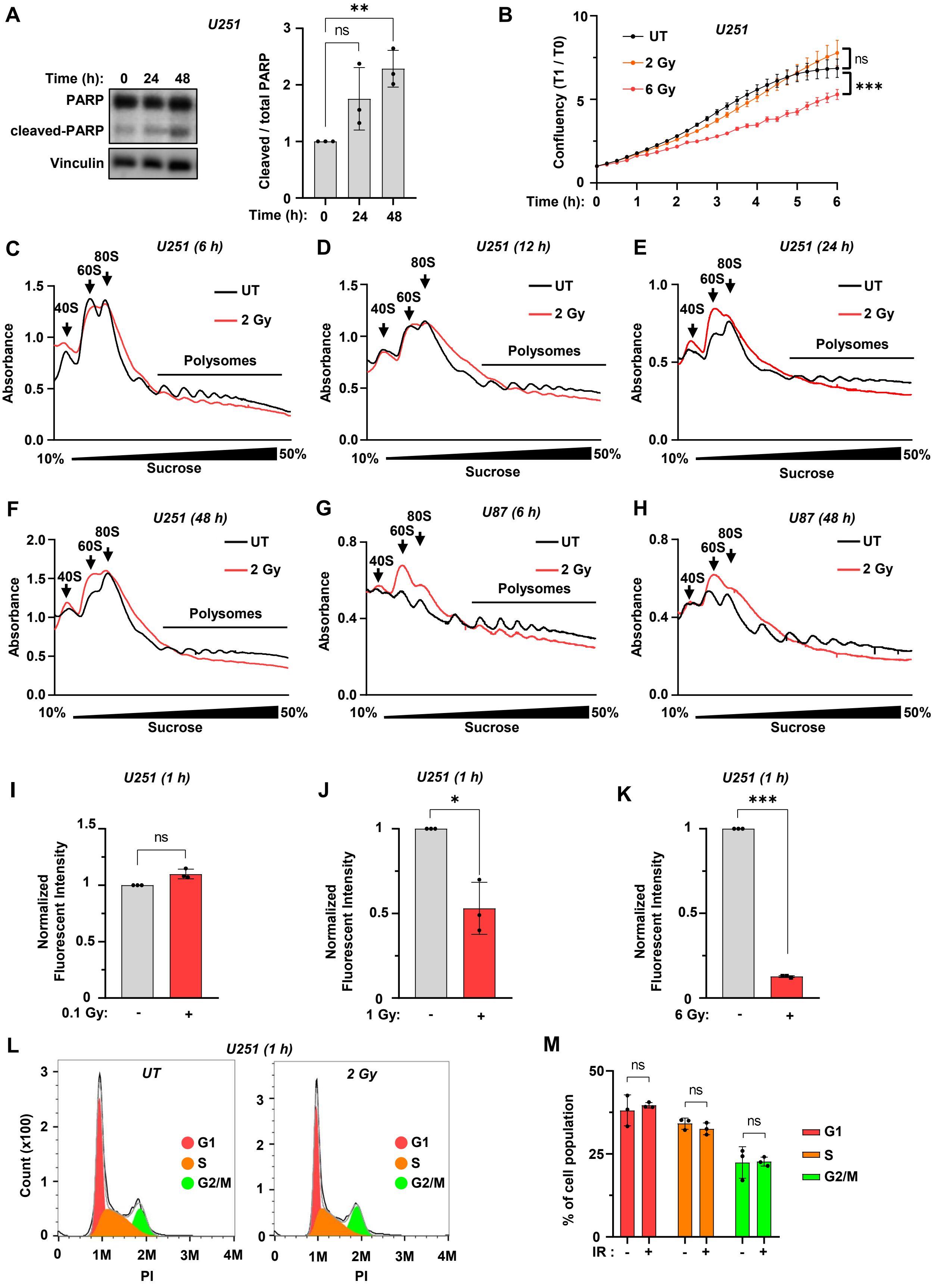


**Supplementary Figure 1: Ionizing radiation represses mRNA translation in GBM cells.** (**A**) *Left:* Western blot analysis of expression of cleaved PARP (c-PARP) using lysates derived from U251 cells 24 h and 48 h after 2 Gy irradiation. Vinculin was used as a loading control. *Right:* Densitometric quantitation of the cleaved-PARP/total-PARP signals normalized to Vinculin for each replicate and presented as mean ±SD; n=3; **$p<0.01$, one-way Anova. (**B**) Incucyte SX1 live-cell analysis of proliferation curves derived from % confluency normalised to starting % confluency after 2 Gy or 6 Gy irradiation of U251 cells. Data are presented as mean ± SD; n=3, two-way Anova; ns=non-significant, ***$p<0.001$. (**C-H**) Representative polysome profiling traces from U251 (C-F) and U87 cells (G & H) at the indicated time points following 2 Gy irradiation. (**I-K**) Quantification of nascent protein synthesis measured by metabolic pulse labelling with the methionine analogue L-homopropargylglycine (HPG) 1 h after irradiation with in 0.1 Gy (I), 1 Gy (J), and 6 Gy (K). Normalized fluorescence intensity was calculated by deduction of the unstained fluorescence followed by division by untreated fluorescence. Data are presented as mean ± SD; n=3, two-way Anova; ns=non-significant, *$p<0.05$, ***$p<0.001$. (**L**) Representative DNA content histograms of U251 cells analyzed by PI staining in U251 cells 1 h following 2 Gy irradiation. (**M**) Quantification of cell cycle distribution in (L). Data are presented as mean ± SD; n=3, one-way Anova; ns=non-significant.

### Supplementary Figures

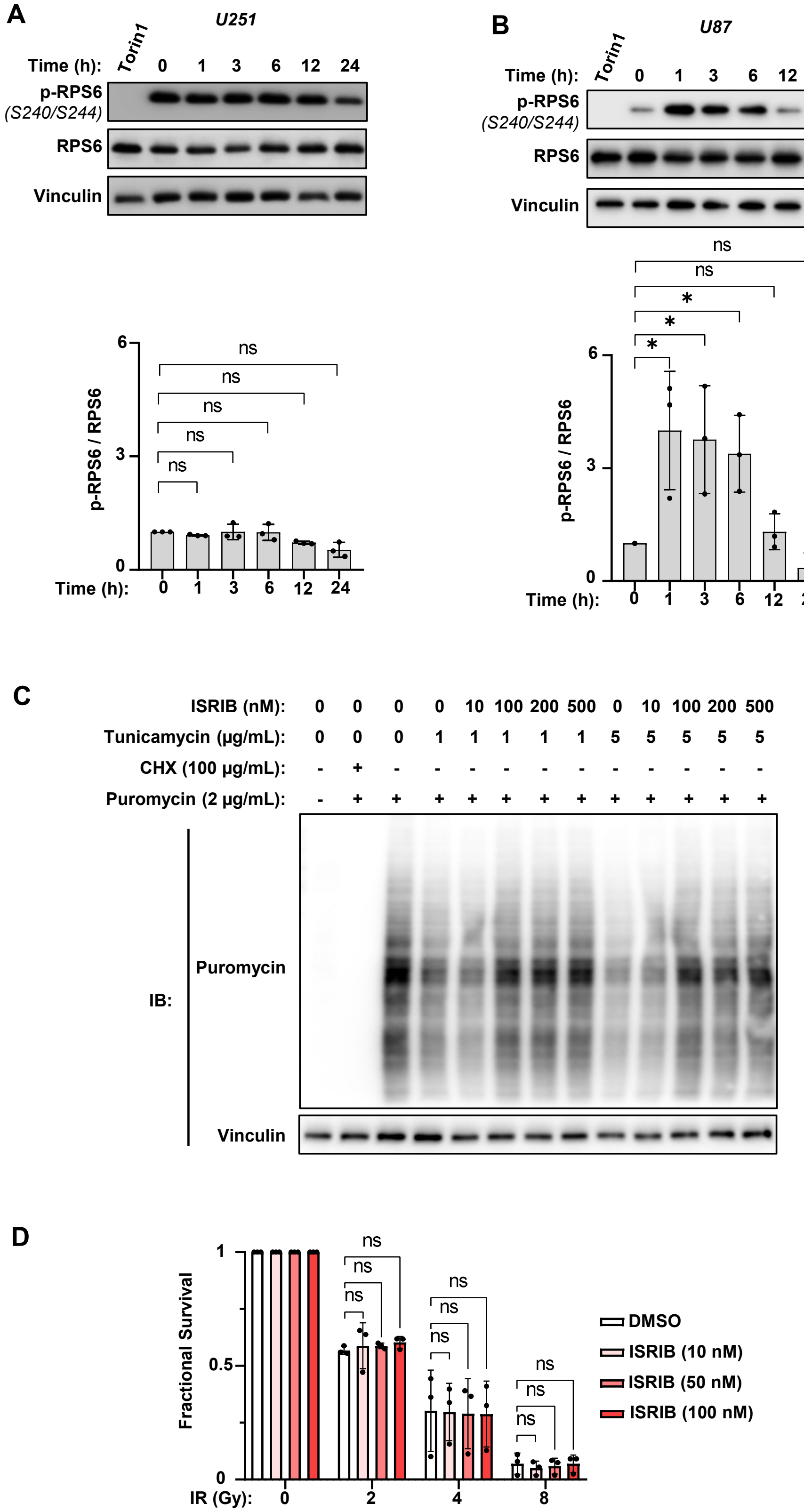


**Supplementary Figure 2: mTOR and ISR-independent cellular response to IR in GBM cells.** (**A & B**) *Top:* Western blot analysis of phosphorylation of RPS6 (Serine 240/244) in U251 (A) and U87 cells (B) following 2 Gy irradiation. Torin-1 (15 nM) treatment for 24 h was used as a control. Vinculin was used as a loading control. *Bottom:* Densitometry quantifications of p-RPS6/RPS6 in the respective cells. (**C**) Quantification of new protein synthesis by Surface Sensing of Translation (SUnSET) assay in U251 cells pre-treated with ISRIB or DMSO for 1 h prior to tunicamycin treatment with indicated dosages for 4 h. CHX treatment was used as a control. Cells were collected following a 15 min pulse with 2 µg/mL puromycin. Vinculin was used as a loading control. (**D**) Colony formation

### Supplementary Figures

assay with U251 cells. Cells were pre-treated with the indicated dose of ISRIB for 1 h followed by 2 Gy irradiation. 10 days post-seeding cells were fixed and fractional survival normalized to plating efficiency. Data are presented as mean ± SD; n=3, one-way Anova; ns=non-significant, *p<0.05.

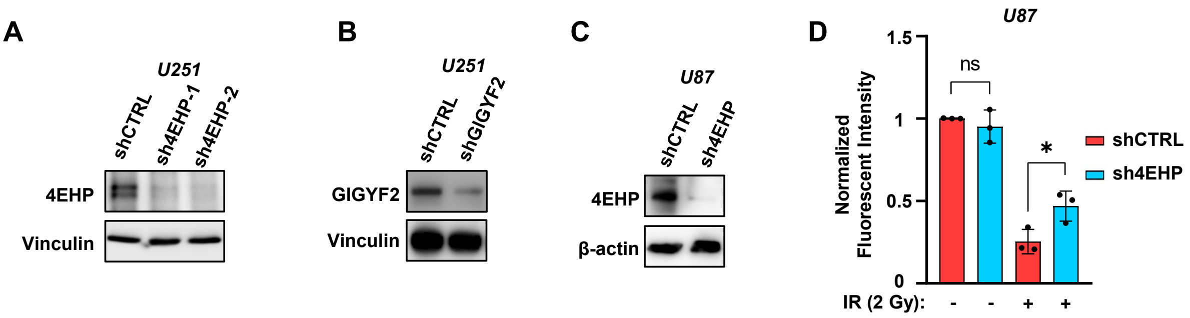


**Supplementary Figure 3. Depletion of 4EHP and GIGYF2 reverses the translational repression upon IR exposure.** (**A**) Western blot analysis of expression of 4EHP in U251 cells with stable expression of two independent shRNAs against 4EHP (sh4EHP-1 and sh4EHP-2) or a control shRNA (shCTRL). (**B**) Western blot analysis of expression of GIGYF2 in U251 cells with stable expression of a shRNA against GIGYF2 (shGIGYF2) or shCTRL. (**C**) Western blot analysis of expression of 4EHP in U87 cells with stable expression of sh4EHP-1 or shCTRL. (**D**) Quantification of nascent protein synthesis in shCTRL and sh4EHP U87 cells measured by metabolic pulse labelling with the methionine analogue L-homopropargylglycine (HPG), 1 h post-treatment with 2 Gy radiation. Signals were normalized against the unstained samples for each group. Data are presented as mean ± SD; n=3, one-way Anova; ns=non-significant, *p<0.05.

### Supplementary Figures

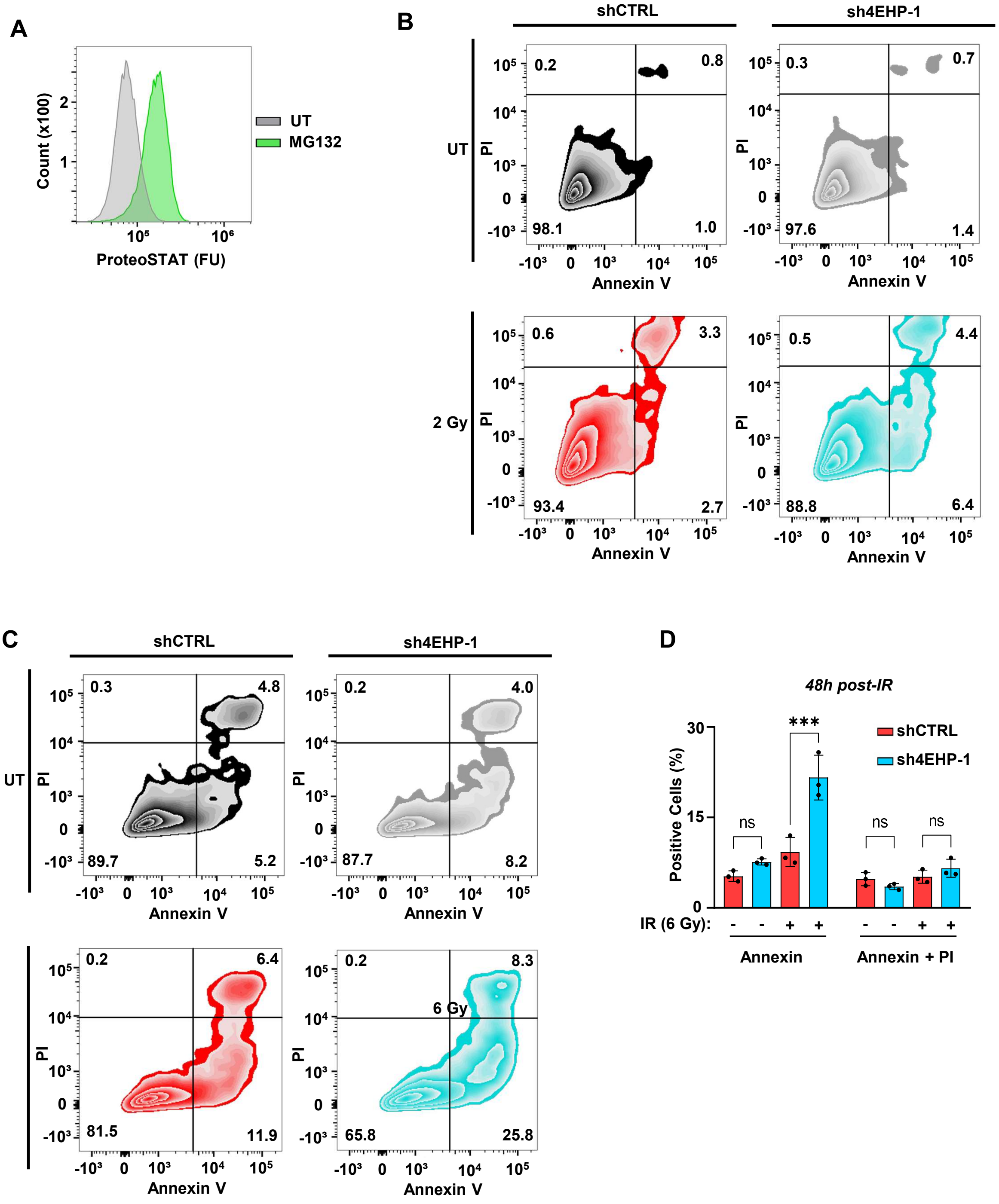


**Supplementary Figure 4. 4EHP contributes to the maintenance of cellular fitness upon IR exposure.** (**A**) Analysis of cellular protein aggregation using ProteoSTAT assay. U251 cells were treated 5 μM of the proteasome inhibitor MG132 for 16 h. ProteoSTAT-stained cells were analysed by flow cytometry. (**B**) Representative flow cytometry density plots from Annexin V/PI stained shCTRL and sh4EHP-1 U251 cells 48 h after treatment with 2 Gy IR. (**C**) Representative flow cytometry density plots from Annexin V/PI stained shCTRL and sh4EHP-1 U251 cells 48 h after treatment with 6 Gy IR. (**D**) Quantification of % of Annexin V-positive or Annexin V+PI-positive shCTRL and sh4EHP-1 U251 cells 48 h after treatment with 6 Gy IR.

### Supplementary Figures

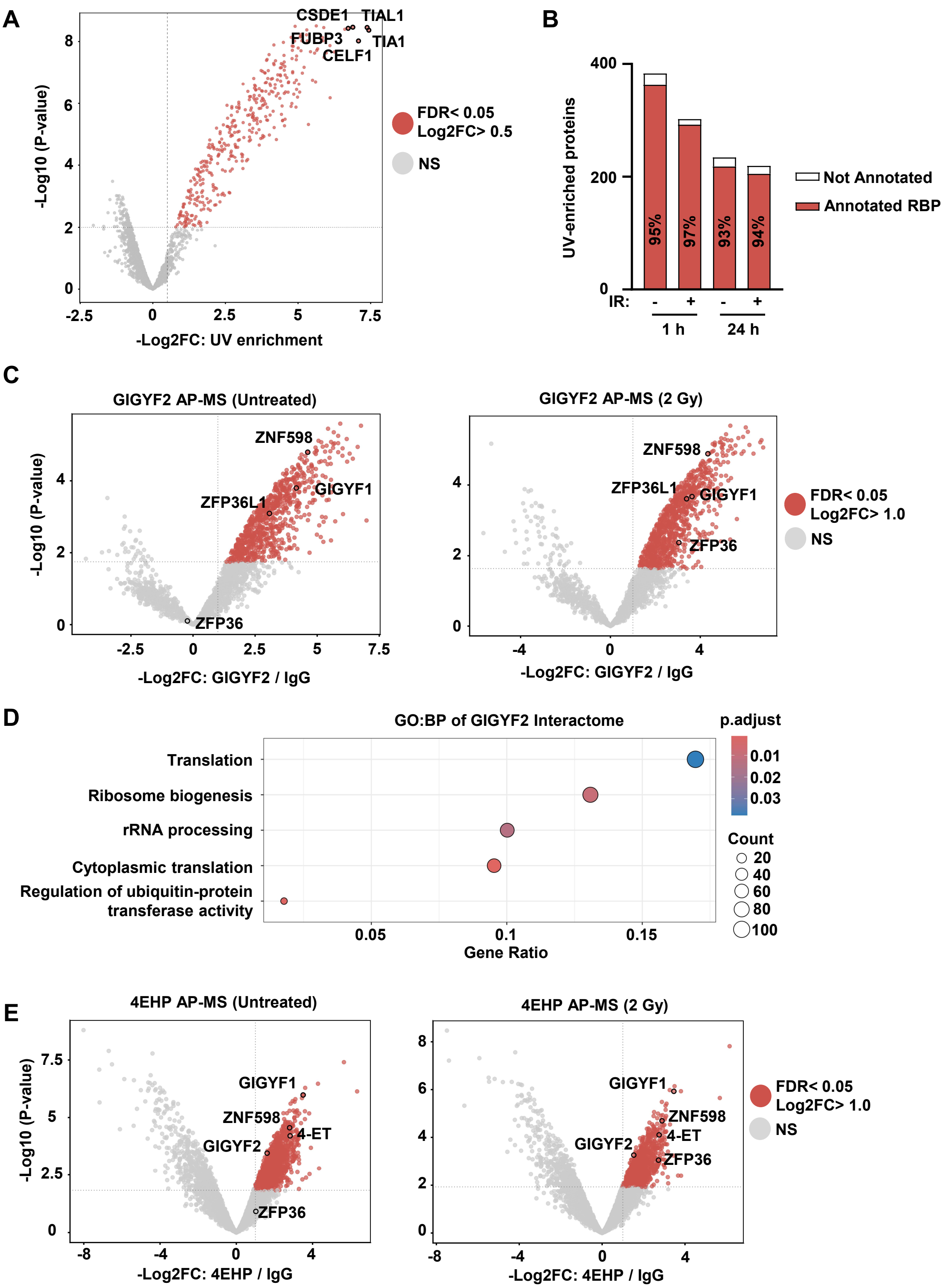


**Supplementary Figure 5. IR-induced changes in RBPome and GIGYF2 or 4EHP protein interactomes in U251 cells.** (**A**) Volcano plot showing the Log2 fold-change and the p-value for protein enrichment by UV crosslinking in the untreated U251 cells. n= 6 independent replicates. Proteins with FDR<0.05 and Log2FC>0.5 are coloured red. (**B**) Bar chart of the proportion of UV-enriched proteins (FDR<0. and Log2FC>0.5) across conditions that were identified as RBPs in ≥3 studies collated in RBPbase database. (**C**) Volcano plots showing the Log2 fold-change and the p-value for protein enrichment by endogenous GIGYF2 immunoprecipitation over IgG control in untreated U251 cells (*Left*), or 1 h after 2 Gy irradiation (*Right*). Proteins with FDR<0.05 are coloured in red. n=3 independent replicates. (**D**) Gene ontology analysis of biological pathways related to proteins significantly enriched (FDR<0.05) by GIGYF2 immunoprecipitation over IgG controls. (**E**) Volcano plots showing the Log2 fold-change and the p-value for protein enrichment by endogenous

**Supplementary Figures**

4EHP immunoprecipitation over IgG control in untreated U251 cells (*Left*), or 1 h after 2 Gy irradiation (*Right*). Proteins with FDR<0.05 are coloured in red. n=3 independent replicates.

### Supplementary Figures

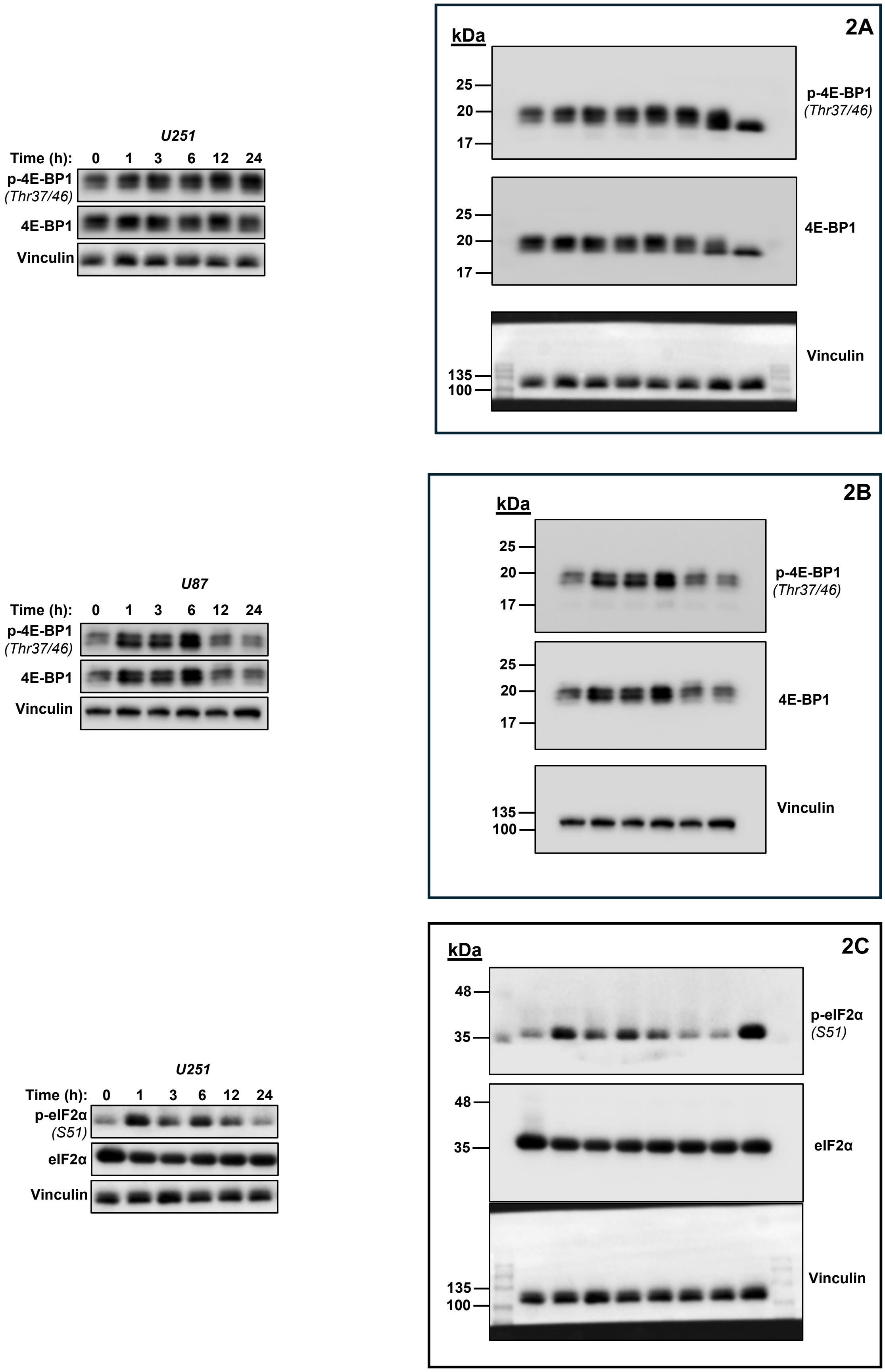


**Supplementary Figures 6:** Uncropped images of blots used in Figure 2A-C.

### Supplementary Figures

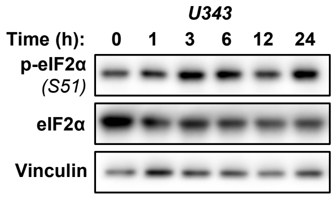


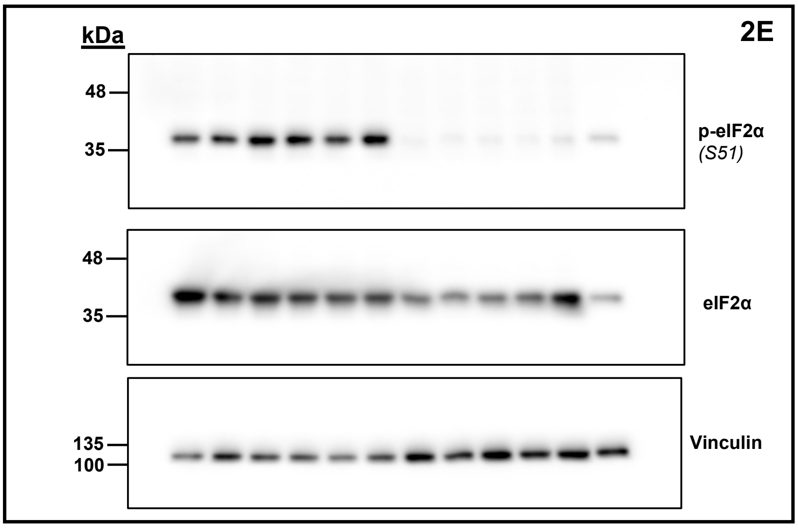


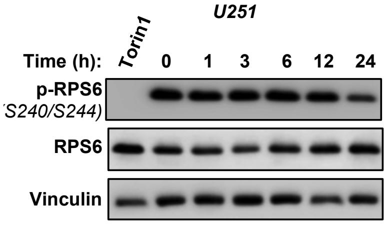


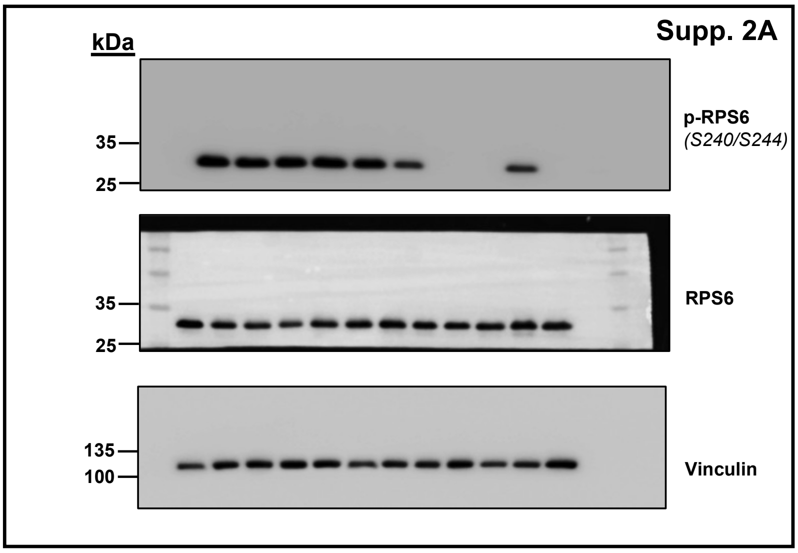


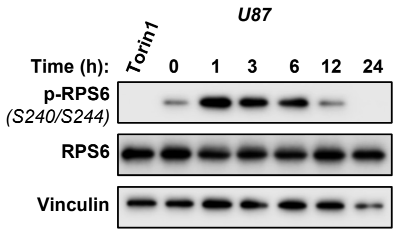


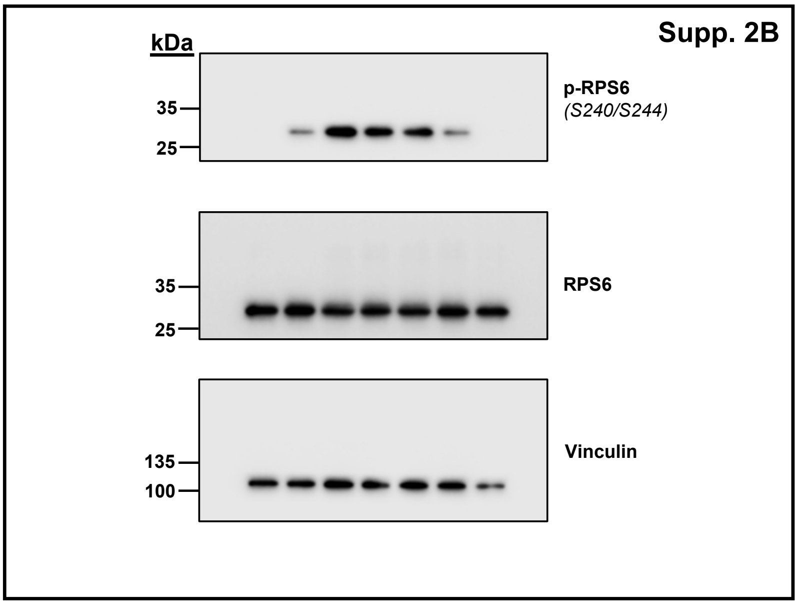


**Supplementary Figures 7:** Uncropped images of blots used in Figure 2 and Supp Fig. 2A & B.

### Supplementary Figures

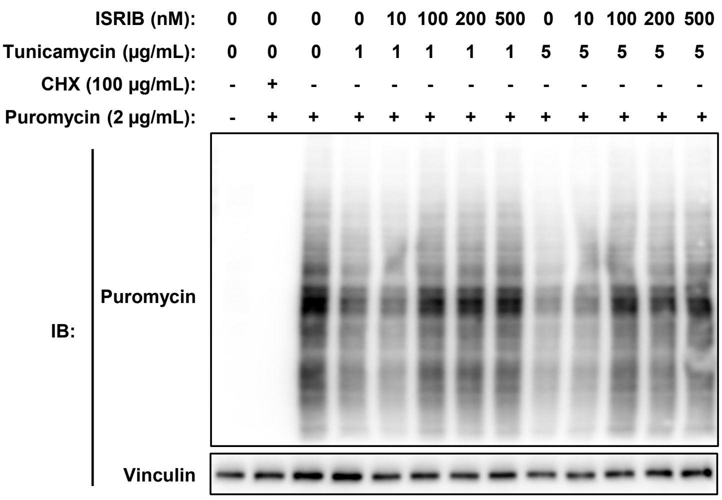


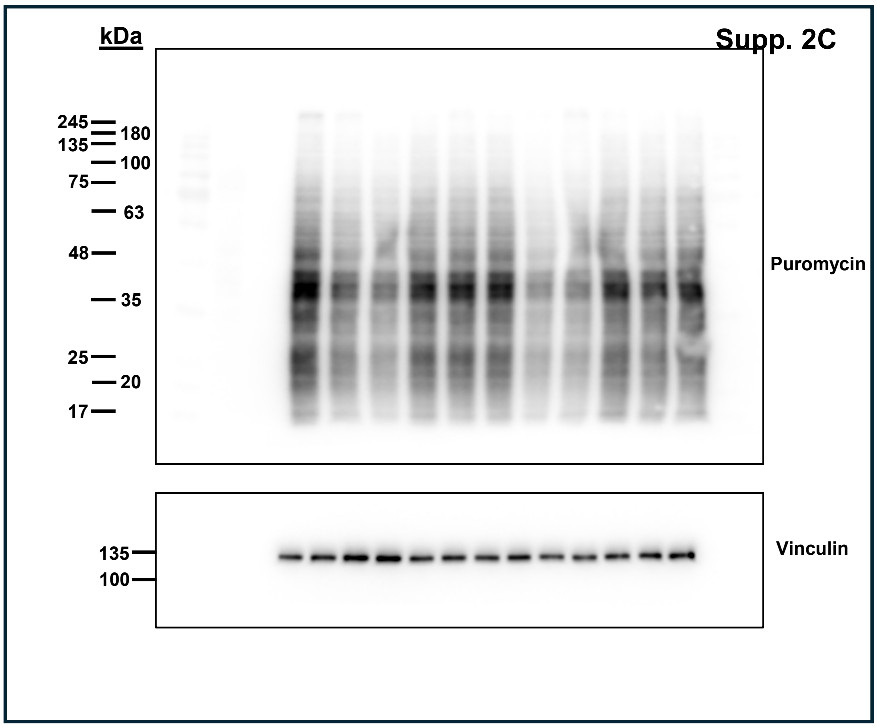


**Supplementary Figures 8:** Uncropped images of blots used in Supp. Figure 2C.

### Supplementary Figures

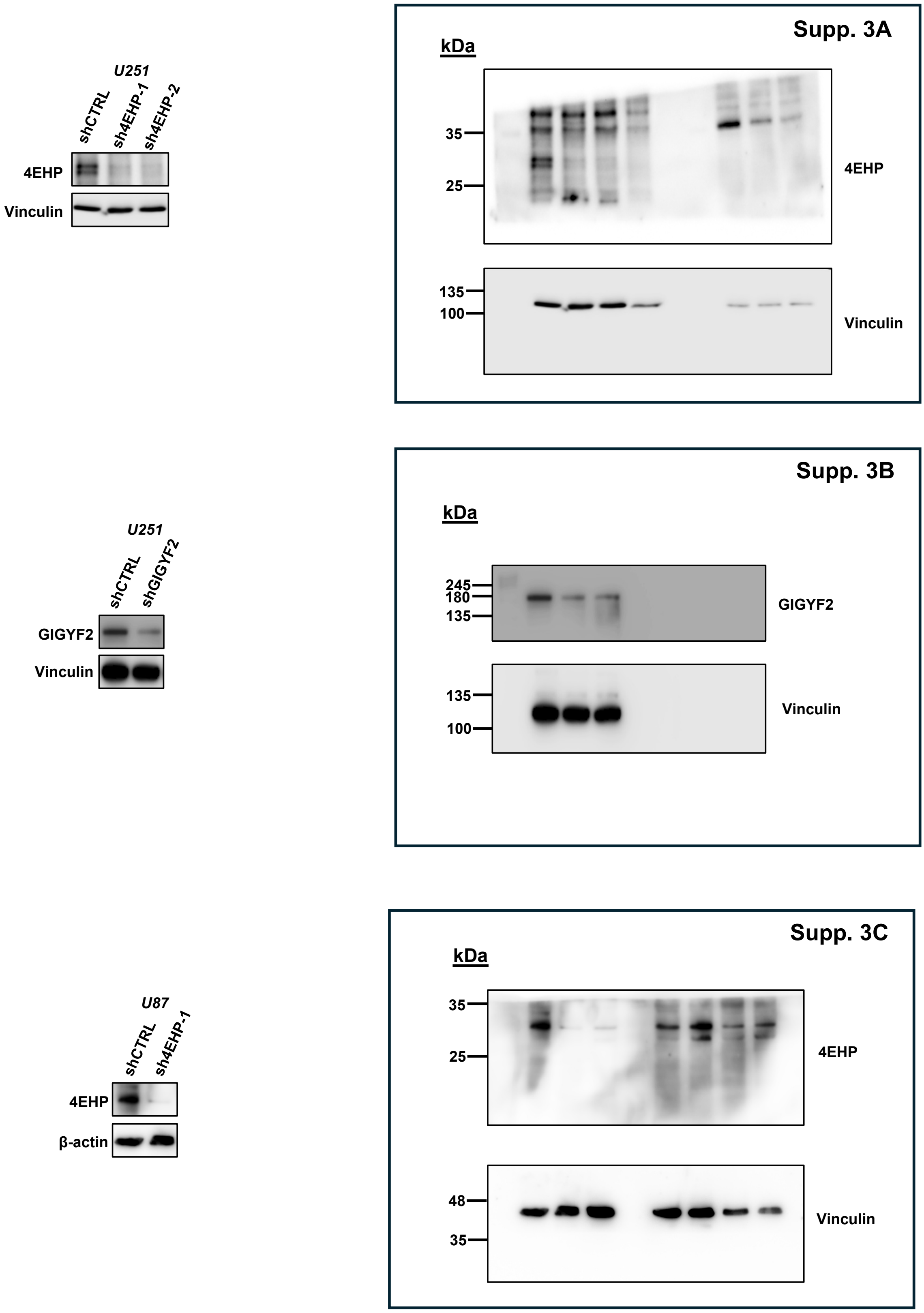


**Supplementary Figures 9:** Uncropped images of blots used in Supp. Figure 3A-C.

**Supplementary Figures**

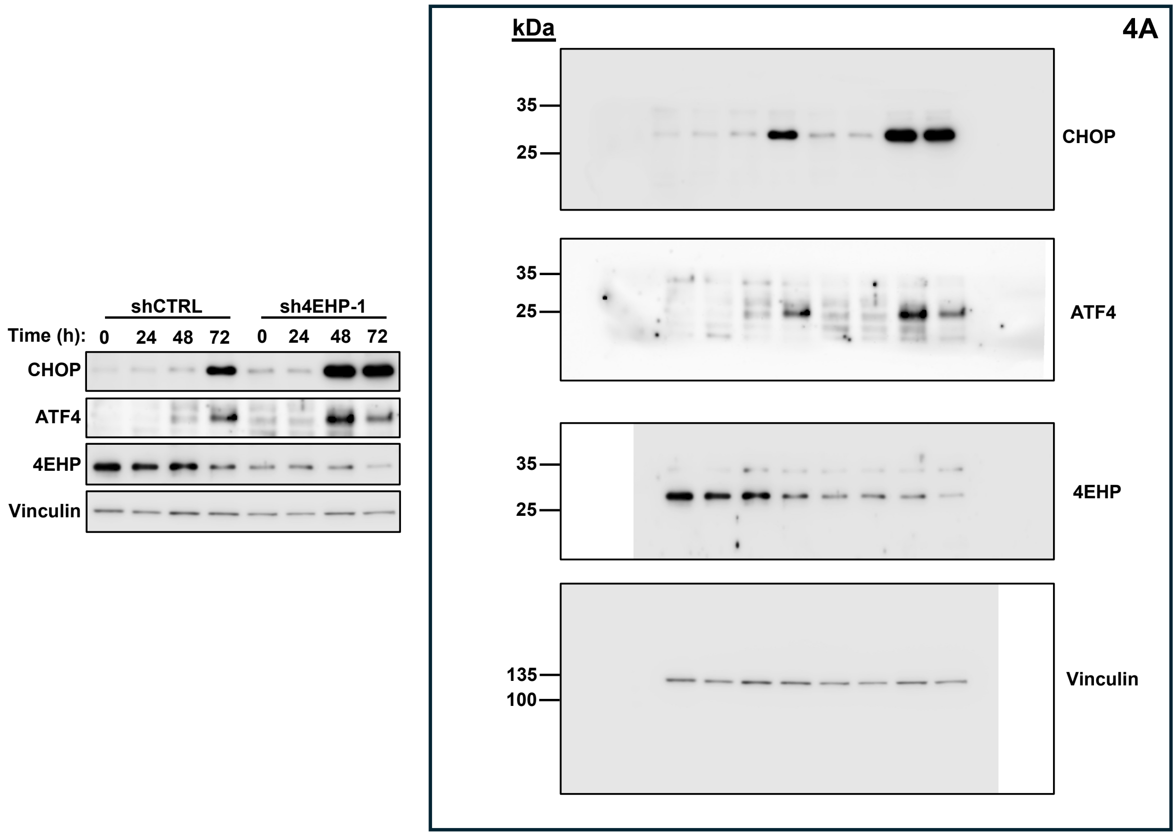


**Supplementary Figures 10:** Uncropped images of blots used in Figure 4A.

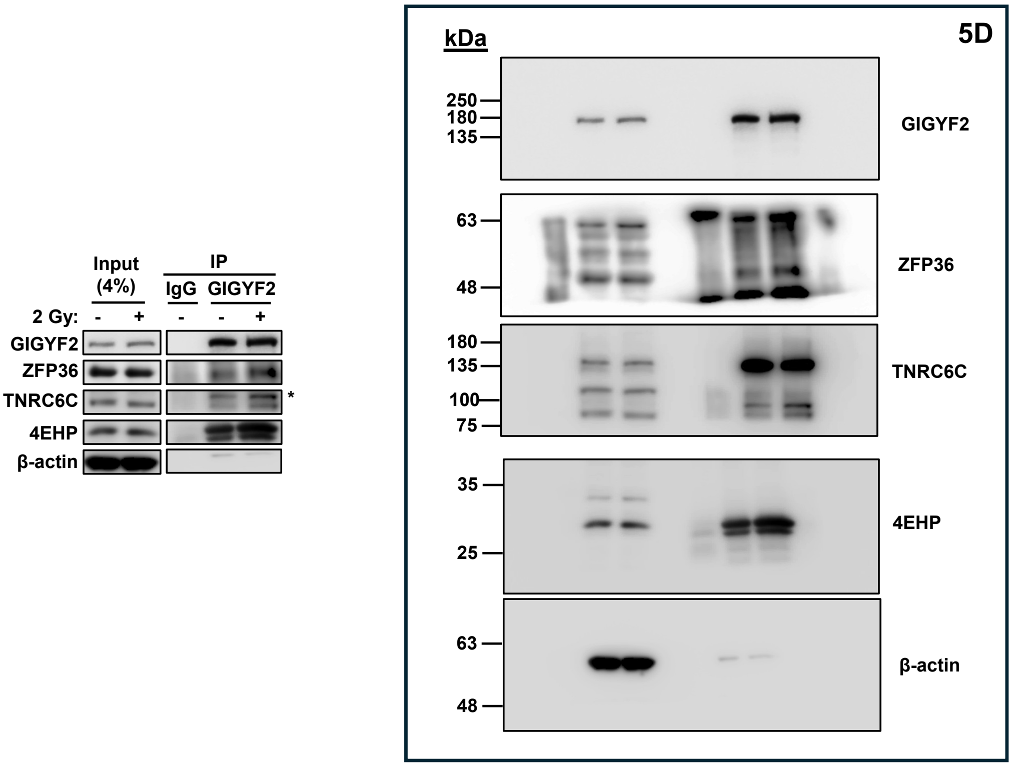


**Supplementary Figures 11:** Uncropped images of blots used in Figure 5D.

### Supplementary Figures

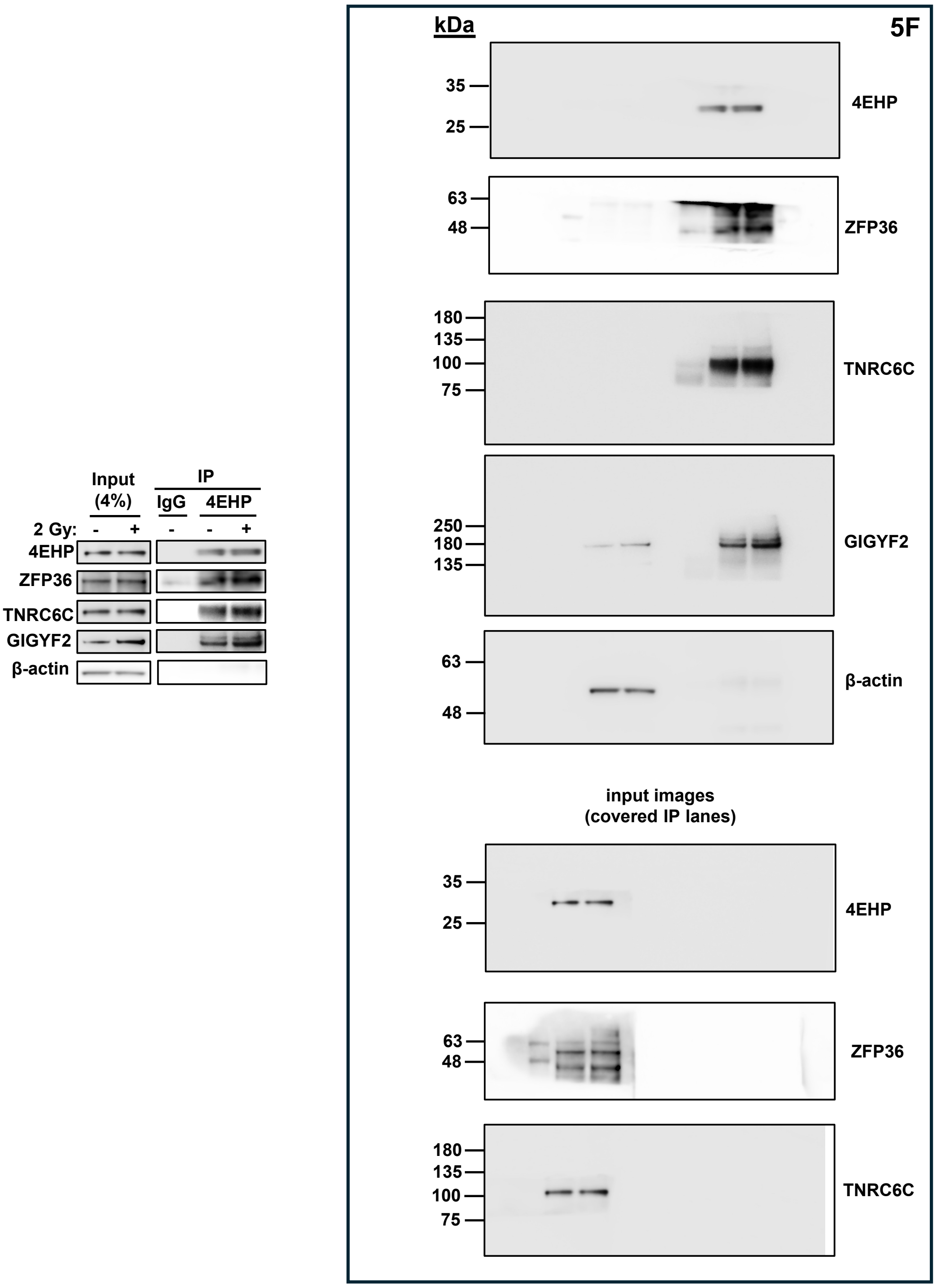


**Supplementary Figures 12:** Uncropped images of blots used in Figure 5F.
